# A tectal reservoir implements adaptive visuomotor transformation via serotonergically coordinated push-pull-like mechanisms

**DOI:** 10.1101/2025.06.10.658856

**Authors:** Yu Qian, Sha Li, Ming-Quan Chen, Hong-Li Wang, Ting-Ting Zhao, Xu-Fei Du, Jiu-Lin Du

**Author notes:** These authors contributed equally (Yu Qian, Sha Li). These authors jointly supervised this work: Jiu-Lin Du, Xu-Fei Du, Yu Qian.

## Abstract

Adaptive visuomotor transformation requires sensorimotor circuits to generate accurate, robust, and flexible outputs, but its underlying cellular mechanism remains poorly understood. Leveraging a zebrafish mesoscopic connectome, we built a biologically constrained spiking neural network of the optic tectum (OT), a conserved vertebrate center for visuomotor transformation, and drove the model with real retinal inputs. Integrating *in silico* and biological interrogations, we revealed that the OT functions as a biologically structured reservoir complemented by serotonergic systems to implement adaptive visuomotor transformation. Within the tectal reservoir, inhibitory interneurons with layer-matched axonal arborizations suppress task-unrelated pathways to ensure visuomotor accuracy, while excitatory interneurons with deep-layer terminating axons reinforce task-related pathways to enhance noise robustness. Furthermore, visually responsive OT-projecting serotonergic subsystems reweight competing pathways to confer visuomotor flexibility. Thus, our findings delineate how specific interneuron-type-embodied push-pull-like mechanisms coordinate with serotonergic neuromodulation to drive adaptive visuomotor transformation, offering a mechanistic framework for neural computational architectures.

## INTRODUCTION

Evolutionarily sculpted biological neural networks can transform sensory inputs into survival-critical behaviors with minimal experience, reflecting innate circuit organization and rapid learning.^1,2^ In contrast, artificial neural networks often rely on extensive data-driven training and can exhibit limited generalization under noise or changing conditions, motivating the search for neural circuit principles that support robust, flexible behavior.^1,3,4^ Understanding how hardwired circuit architectures achieve effective sensorimotor processing remains fundamental to deciphering biological intelligence.^5^ Vision serves as a primary sensory modality for interpreting dynamic environments. The superior colliculus and its homolog, the optic tectum (OT), constitute an evolutionarily conserved visuomotor transformation center in vertebrates^6–8^ that converts retinotopic visual inputs into signals initiating context-appropriate behaviors, such as orienting and escape.^8–11^ While the afferent and efferent pathways of the OT have been extensively characterized,^7,12–16^ its intrinsic circuits remain less well defined, especially how distinct neuronal types coordinate to implement adaptive visuomotor transformation.

The OT comprises a sparse and heterogeneous recurrent network formed by diverse tectal interneurons (TINs) and tectal projection neurons (TPNs).^6,17,18^ This architecture is inherently suited for transforming sensory inputs into distributed, time-evolving activity patterns. Within reservoir-computing frameworks, such recurrent network can serve as a dynamic substrate from which downstream readouts can extract task-relevant outputs without task-specific rewiring.^19,20^ Meanwhile, neuromodulatory signals can reconfigure circuit states without changing anatomical wiring, allowing the same recurrent architecture to support context-dependent computations.^21–23^ This prompted us to ask whether stable OT wiring generates a tectal activity reservoir, and whether OT-innervating neuromodulatory systems flexibly reshape its pathway-specific outputs during adaptive visuomotor transformation.

Larval zebrafish (*Danio rerio*) serves as a tractable vertebrate model for dissecting mechanisms of adaptive visuomotor transformation, owing to its conserved tectal architecture,^6,17^ optical transparency for cellular-resolution activity mapping,^24^ and the availability of the single-cell morphology-based mesoscopic connectome at 6 days post-fertilization (dpf).^18^ However, the marked heterogeneity of OT neurons, coupled with limited genetic access to many of these cell types, hinders the systematic causal dissection of neuron type-specific functions in visuomotor transformation.^6,17,18^ Therefore, we constructed a biologically constrained spiking neural network (SNN) model of the OT, incorporating retinal ganglion cells (RGCs) as input neurons, TINs as an intermediate layer, and TPNs as output neurons. The model was grounded in the zebrafish mesoscopic connectome^18^ and driven by visually evoked activity of RGCs. Unlike abstract models, this model preserves biological realism by incorporating neuronal diversity, anatomically structured connectivity, and experimentally measured inputs, thereby enabling cell-type-resolved *in silico* perturbations that are currently difficult to implement in biological experiments.^25–28^

Leveraging this model, we recapitulated the biological phenomenon that anatomically distinct visuomotor pathways, constituted by RGCs and TPNs, generate pathway-specific readouts. By combining morphotype-resolved *in silico* perturbations with biological experiments, we revealed that distinct TIN morphotypes underpin the accuracy and robustness of visuomotor transformation through a push-pull-like mechanism: specific inhibitory TINs enhance accuracy by suppressing task-unrelated pathways (“push”), while excitatory TINs promote robustness by amplifying task-related pathway responses (“pull”). Moreover, pretectal serotonergic (5-HT) inputs to the OT bias competition between visuomotor pathways by directly modulating TPN responses, thereby conferring visuomotor flexibility as well as enhancing push-pull effects. Collectively, our study elucidates how structured connectivity and neuromodulatory modulation coordinatedly implement adaptive computation of the OT.

### Anatomically segregated tectofugal pathways mediating distinct visuomotor responses

The zebrafish OT comprises morphologically diverse types of tectal neurons (TNs) that mediate visuomotor transformation.^6,17^ To dissect its complex architecture, we analyzed the reconstructed single-cell morphologies of 1,904 TNs within the larval zebrafish mesoscopic atlas.^18^ The reconstructed TNs included 718 TPNs and 1186 TINs (Figures 1A and S1A-S1D). TPNs, the OT’s output neurons with axons projecting to downstream regions, included 658 excitatory (e-TPNs) and 60 inhibitory neurons (i-TPNs) (Figures S1A and S1B). TINs, local interneurons with axons confined within the OT, comprised 286 excitatory (e-TINs) and 900 inhibitory neurons (i-TINs) (Figures S1C and S1D).

**Figure 1.**
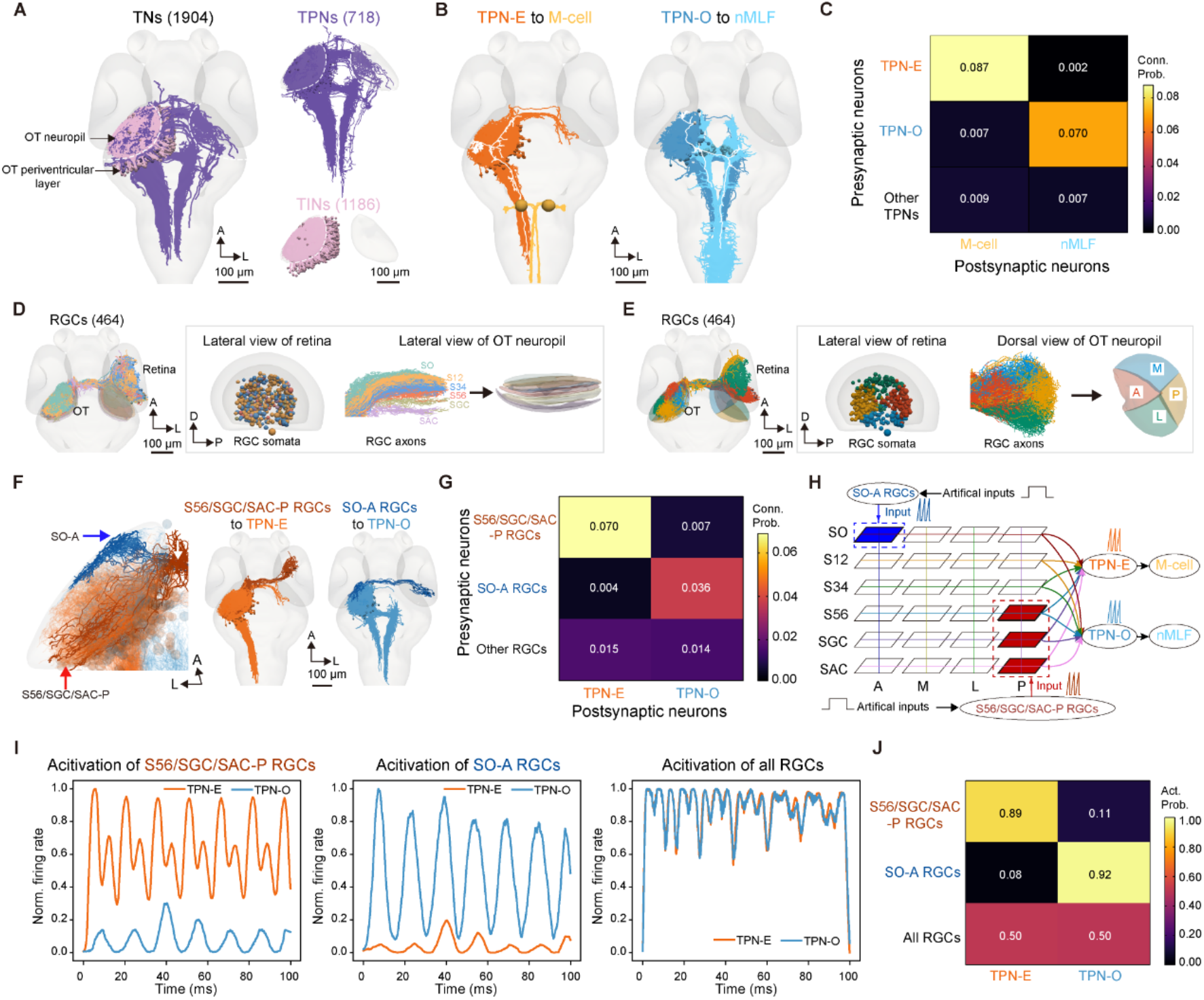
Mesoscopic connectome-constrained SNN modeling of two distinct visuomotor pathways in the zebrafish optic tectum. (A) Reconstructed single-cell morphologies of tectal neurons (TNs) from the larval zebrafish mesoscopic atlas^18^, classified into tectal projection neurons (TPNs, purple) with axons exiting the optic tectum (OT), and tectal interneurons (TINs, pink) with axons confined within the OT. Dashed line denotes the OT soma-neuropil boundary. Neurons were mirrored to the left hemisphere for visualization. Numbers in parentheses indicate the number of reconstructed neurons. (B) Two distinct tectofugal pathways are defined by the putative connectivity between TPN axons and downstream neurons’ dendrites. Left, escape pathway constituted by escape-related TPNs (TPN-E, orange) and their downstream Mauthner cells (M-cells, yellow). Right, orienting pathway constituted by orienting-related TPNs (TPN-O, blue) and their downstream neurons in the nucleus of the medial longitudinal fasciculus (nMLF, light blue). White traces are representative morphologies. (C) Putative connection probability between TPN subtypes and downstream targets (inferred from axon-dendrite appositions; see STAR Methods). The numbers indicate connection probabilities. (D) Laminar organization of the OT neuropil defined by axonal terminations of reconstructed retinal ganglion cells (RGCs). RGCs are color-coded by the OT neuropil stratum of their axonal termination: stratum opticum (SO), stratum fibrosum et griseum superficiale 1 - 2 (SFGS1-2; S12), SFGS3 - 4 (S34), SFGS5 - 6 (S56), stratum griseum centrale (SGC), and stratum album centrale (SAC). (E) Retinotopic organization of the OT neuropil inferred from RGCs soma positions: anterior (A), posterior (P), medial (M), and lateral (L). RGCs are color-coded by soma position in the retina. (F) Putative pathways from RGCs to TPN-E and TPN-O. Left, representatives of RGCs projecting to posterior S56, SGC, or SAC (“S56/SGC/SAC-P RGCs”, brown) and RGCs projecting to anterior SO (“SO-A RGCs”, dark blue). Middle putative connectivity from S56/SGC/SAC-P RGCs to TPN-E. Right, putative connectivity from SO-A RGCs to TPN-O. The white arrow indicates RGC axons before entering the OT neuropil; the blue arrow indicates the neuropil SO-A microdomain; the red arrow indicates the S56/SGC/SAC-P microdomains. (G) Putative connection probabilities from RGC populations to TPN-E and TPN-O. The numbers indicate connection probabilities. (H) Schematic of the mesoscopic connectome-constrained SNN model of RGC-TPN pathways. Selective activation of S56/SGC/SAC-P or SO-A RGC populations was implemented by applying a 100-ms constant input step to the corresponding RGCs. (I) TPN responses evoked by selective activation of RGC populations *in silico*. Left, Stimulation of S56/SGC/SAC-P RGCs preferentially drives TPN-E. Middle Stimulation of SO-A RGCs preferentially drives TPN-O. Right, stimulation of all RGCs co-activates both pathways. Traces show the mean across 5 simulation trials. (J) Activation probability of TPN-E and TPN-O under different RGC populations stimulation conditions. The numbers indicate activation probabilities. A, anterior; P, posterior; D, dorsal; L, lateral; M, medial. See also Figures S1 and S2.

The transformation of external visual scenes into behaviorally relevant motor commands necessitates selective routing of sensory signals to specific motor circuits, a process requiring precise topological alignment between input and output structures. To elucidate how the OT achieves this routing fidelity, we focused on two subsets of e-TPNs defined by their target brain regions: escape-related TPNs (TPN-E), which connect to Mauthner cells (M-cells) to drive rapid escape behaviors,^29–32^ and orienting-related TPNs (TPN-O), which connect to neurons in the nucleus of the medial longitudinal fasciculus (nMLF) to mediate orienting and postural control.^12,33–35^ Using the zebrafish mesoscopic atlas and the iBrAVE analysis platform,^18,36^ we quantified the putative connections between TPNs and M-cells or nMLF neurons based on axon-dendrite proximity in the registered common space (see STAR Methods), identifying 223 TPN-E (209 with ipsilateral and 14 with bilateral processes in the OT neuropil), and 62 TPN-O (27 with ipsilateral, 33 with contralateral, and 2 with bilateral projections) (Figures 1B, 1C, and S1E). Beyond their defining downstream targets, TPN-E and TPN-O exhibited significantly different projection profiles across a broad range of downstream brain regions (Figure S1F), suggesting that they recruit distinct downstream modules for orchestrating behaviors. In addition, beyond TPN-O and TPN-E, the remaining e-TPNs can be further subdivided into different subtypes based on their soma location and downstream target regions (Figure S1G). This output-specific parcellation reflects a conserved topographic organization,^6,12^ wherein anatomically segregated tectofugal pathways process behaviorally distinct visuomotor streams.

### Anatomically segregated RGC inputs to distinct tectofugal pathways

To resolve visual afferents of TPN-E and TPN-O tectofugal pathways, we analyzed the reconstructed single-cell morphologies of 464 RGCs within the zebrafish mesoscopic atlas.^18^ The majority of RGCs (461/464) sent axon terminals to the neuropil of contralateral OT and formed a retinotopic projection map (Figure S2A). Individual RGC axons exclusively arborized within one of six distinct OT neuropil strata, arranged from superficial to deep: stratum opticum (SO), stratum fibrosum et griseum superficiale 1-2 (SFGS1-2, S12), SFGS3-4 (S34), SFGS5-6 (S56), stratum griseum centrale (SGC), and stratum album centrale (SAC, also referred to as SAC/SPV, denoting the neuropil-periventricular boundary) (Figures 1D and S2B). Consistent with retinotopic organization principles,^6,7,17^ RGC soma positions well predicted axonal termination sectors within four topographic domains: anterior (A), posterior (P), medial (M), and lateral (L) (Figures 1E and S2C). Combining these two axes yielded 24 lamina-by-topography microdomains for visuomotor processing. We therefore grouped RGCs into 24 populations based on the lamina and topographic sector of their OT arborizations.

Before terminating in the OT neuropil, some RGCs (272/461) also sent collateral arbors in distinct 9 arborization fields (AFs) (Figure S2D), which are known to encode different visual features.^37^ Consistent with previous findings, we found a functional association between the AF and OT neuropil targeting of RGC axonal arborizations. In particular, RGCs with collaterals in AF7, a region implicated in prey detection,^38,39^ formed dense axonal terminations in the anterior or lateral SO. In contrast, RGCs with collaterals in AF9, which encodes global luminance and large-object stimuli,^29,39,40^ predominantly targeted the posterior or anterior SGC/SAC with minor projections to S56 (Figure S2D). Given that anterior superficial layers (SO-A) of the OT neuropil process prey-related cues, while posterior deep OT domains (centered on SGC/SAC, with minor contribution from posterior S56) are preferentially engaged by predator-related or large-object visual cues,^17,29,37,41^ we analyzed putative connections between these RGC axons and TPN dendrites. We found that S56/SGC/SAC-P- and SO-A-projecting RGCs preferentially connected with TPN-E and TPN-O, respectively (Figures 1F and 1G). In contrast, RGCs targeting other OT microdomains showed no connection preference between TPN-E and TPN-O, exhibiting relatively low connection probabilities for both (Figure 1G).

These results demonstrate two anatomically segregated visuomotor pathways: an escape-associated pathway receiving predominant retinal input from posterior deep OT domains (S56/SGC/SAC-P), converging onto TPN-E, and driving motor responses through M-cells; and an orienting pathway initiating in the SO-A region, connecting to TPN-O, and triggering behavior via nMLF neurons.

### SNN modeling of RGC-TPN pathways

To model RGC-TPN pathways, we constructed a biologically constrained SNN that incorporated reconstructed connectivity between RGCs and TPNs (Figure 1H). The model used leaky integrate-and-fire (LIF) neurons parameterized with zebrafish neuronal properties (Table S1; see STAR Methods). Selective activation of S56/SGC/SAC-P RGCs evoked strong spiking in TPN-E but only weakly recruited TPN-O (Figures 1I, left and 1J). Conversely, selective activation of SO-A RGCs robustly engaged TPN-O, with minimal recruitment of TPN-E (Figures 1I, middle and 1J). These simulations indicate that the anatomical segregation is sufficient to generate pathway-biased outputs *in silico*.

Although a visual stimulus typically evokes a specific behavior, it usually engages a broad repertoire of RGCs.^29^ We recapitulated this scenario by globally stimulating all RGCs in the model. Intriguingly, global activation of all RGCs activated both TPN-E and TPN-O (Figures 1I, right and 1J), suggesting the concurrent occurrence of escape and orienting behaviors in the model. However, simultaneous engagement of these two visuomotor pathways is implausible in natural contexts, since escape and orienting are usually mutually exclusive within individual epochs.^42,43^ This implies that additional circuit mechanisms are required to resolve pathway competition under widespread retinal drive.

### A biologically structured TIN reservoir enables pathway-selective visuomotor transformation

Besides TPNs, the OT contains diverse TINs (see Figures 1A, S1C, and S1D) implicated in visual processing.^44–48^ To improve the biological realism of the model, we then constructed a biologically constrained TIN network and incorporated it into the RGC-TPN SNN.

To construct the TIN network, we analyzed the reconstructed single-cell morphologies of 286 e-TINs and 900 i-TINs within the larval zebrafish mesoscopic atlas^18^ (see Figures S1C and S1D). Based on the distinct stratification patterns of their neurites within the OT neuropil, these neurons were classified into 20 morphotypes of e-TINs (Figures S3A-S3C) and 24 morphotypes of i-TINs (Figures S3D-S3F). In line with established TIN classification schemes from previous studies,^45,46^ we categorized these neurons into three canonical arborization patterns for network construction: 1) monostratified TINs, which confine both dendrites and axons to a single lamina of the OT neuropil; 2) multilayered TINs, whose dendrites occupy superficial laminae while their axons arborize in deeper laminae; and 3) non-stratified TINs, which exhibit intermingled dendritic and axonal processes without laminar restriction. The discrimination of TIN axonal and dendritic compartments was achieved by a machine-learning algorithm based on established morphological criteria and molecular labeling of presynaptic sites (Figures S4A-S4C; see STAR Methods). A TIN network was then built based on connection probabilities among all 44 TIN morphotypes, TPN-E, and TPN-O (Figures S4D-S4G and Table S2; see STAR Methods). The TIN network was dominated by inhibitory connections (Figure S4H), and its in-degree distribution was skewed toward i-TINs (Figure S4I). Unlike the Erdős–Rényi (ER) random network and the Barabási-Albert (BA) scale-free network, the TIN network exhibited a unique combination of low clustering, low average path length, high degree assortativity, and low network density, indicating that the TIN network has a topology distinct from the ER and BA baseline networks (Figures S4J-S4N).

To test whether the recurrent TIN network behaves as a reservoir, we analyzed perturbation decay and fading memory (Figures S4O and S4P; see STAR Methods). Under identical RGC inputs, perturbation-induced differences in TIN state trajectories decayed gradually, persisting longer than in the no recurrent control and comparable to the ER and BA reservoirs (Figure S4O). Furthermore, similar to the ER and BA reservoirs, TIN states retained decodable information about recent RGC inputs over a finite delay window (Figure S4P), consistent with the TIN network functioning as a stable, input-driven reservoir.

To drive the model with real biological inputs, we measured RGC activity evoked by one of two ethologically relevant stimuli in 6-8 dpf larvae (Figures 2A and 2B; see STAR Methods): a dark looming stimulus (mimicking an approaching predator) that triggers escape,^29,40^ and a black small moving dot (SMD, 4° in diameter, mimicking a moving prey) that elicits orienting.^39,42^ Both stimuli evoked layer-preferential calcium activity of RGC axonal arbors in the OT neuropil, yet the induced activity spread across the neuropil (Figure 2B, middle), indicating a need for tectal routing mechanisms to process these distributed signals and generate specific visuomotor outputs. The calcium activity served as the model inputs after conversion to spike-proxy current inputs (see STAR Methods).

**Figure 2.**
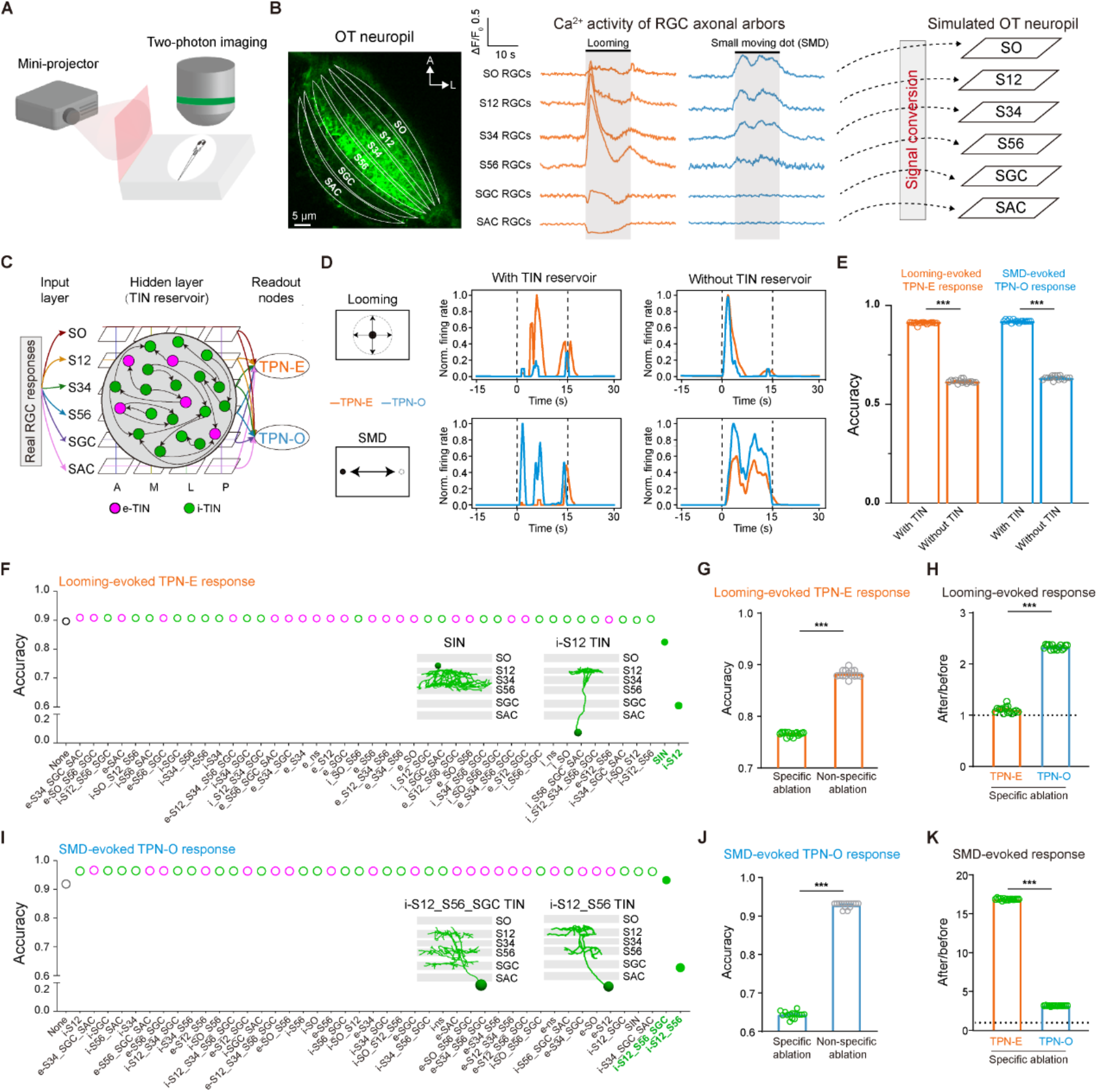
Specific inhibitory TIN morphotypes are required for visuomotor transformation accuracy. (A) Experimental setup for visual stimulation and *in vivo* two-photon calcium imaging in head-fixed larval zebrafish. (B) Layer-specific calcium activity of RGC axonal arbors in the OT neuropil serving as the model inputs. Left, fluorescence image of the OT neuropil (dorsal view; right hemisphere). Middle looming- and small moving dot (SMD)-evoked RGC responses across neuropil laminae. Right, schematic showing RGC activity fed into the model after conversion to spike-proxy current inputs. Activity traces represent the mean ± SEM averaged from 6 larvae (2 trials per larva); error bars are too small to be visible. SO RGCs: SO-projecting RGCs; S12 RGCs: S12-projecting RGCs; S34 RGCs: S34-projecting RGCs; S56 RGCs: S56-projecting RGCs; SGC RGCs: SGC-projecting RGCs; SAC RGCs: SAC-projecting RGCs. The imaging was performed on double-transgenic *Tg(-7atoh7:GAL4-VP16);Tg(UAS:GCaMP6s)* larvae at 6-8 days post-fertilization (dpf), in which RGCs express the calcium indicator GCaMP6s. A SMD of 4° diameter was used. (C) Schematic of OT SNN network architecture. RGC responses serve as inputs (input layer), are processed by the TIN reservoir (hidden layer), and are read out by TPN-E and TPN-O nodes (output layer). (D) Activity of TPN-E (orange) and TPN-O (blue) evoked by looming (top) and SMD (bottom) stimuli with (middle) and without (right) the TIN reservoir. Left, schematic of visual stimuli which last 15 s. Window between two dashed lines indicates the visual stimulation period. The traces are averaged from 5 simulation trials. (E) Response accuracy of looming-evoked TPN-E and SMD-evoked TPN-O responses with or without the TIN reservoir. Looming accuracy was defined as *A*_*E*_/(*A*_*E*_ + *A*_*O*_); SMD accuracy as *A*_*O*_/(*A*_*E*_ + *A*_*O*_), where *A*_*E*_ and *A*_*O*_ are the AUCs of TPN-E and TPN-O responses measured within the stimulation window, respectively. Each small circle on the bars denotes one simulation trial (n = 15 trials). (F) Effects of *in silico* cumulative ablation of 44 TIN morphotypes on the accuracy of looming-evoked TPN-E responses, showing that i-S12 TIN and SIN morphotypes (filled green dots) are the most critical. Inset, typical cases of i-S12 TINs and SINs. e-TINs, magenta circles; i-TINs, green circles. (G) Looming response accuracy after specific ablation of both i-S12 and SIN (“specific ablation”), or ablation of all remaining morphotypes (“Non-specific ablation”). Each circle denotes one simulation trial (n = 15). (H) Specific ablation-induced changes in looming-evoked TPN-E and TPN-O responses. The dashed line indicates the no-change level (fold-change = 1). Each circle denotes one simulation trial (n = 15). (I) Effects of *in silico* TIN cumulative ablation on the accuracy of SMD-evoked TPN-O responses, showing i-S12_S56 and i-S12_S56_SGC TIN morphotypes (filled green dots) are the most critical. Inset: typical cases of i-S12_S56 and i-S12_S56_SGC TINs. e-TINs, magenta circles; i-TINs, green circles. (J) SMD response accuracy after specific ablation of both i-S12_S56 and i-S12_S56_SGC TINs, or non-specific ablation of all remaining morphotypes. Each circle denotes one simulation trial (n = 15). (K) Specific ablation-induced changes in SMD-evoked TPN-E and TPN-O responses. The dashed line indicates the no-change level (fold-change = 1). Each circle denotes one simulation trial (n = 15). Each point represents one independent simulation trial (E and G - K). **\*\*\****P* < 0.001 (two-sided Wilcoxon signed-rank test). See also Figures S3 and S4.

We then integrated the TIN reservoir network as a hidden layer into the RGC-TPN SNN and drove the model with real RGC activity (Figure 2C). The model exhibited visuomotor responses with high accuracy: looming and SMD stimuli selectively activated TPN-E and TPN-O, respectively (Figures 2D, middle and 2E). Importantly, loss of the TIN network caused co-activation of both visuomotor pathways by either stimulus (Figure 2D, right; comparing with Figure 1I, right), leading to reduced accuracy of visuomotor responses for both stimuli (Figure 2E; *P* < 0.001, two-sided Wilcoxon signed-rank test on response accuracy across 15 simulation trials). These results suggest that the TIN network works as a biologically structured reservoir to support pathway-selective visuomotor transformation.

### Specific inhibitory TIN morphotypes ensure visuomotor accuracy by suppressing task-unrelated pathways

To further examine which TIN morphotypes are predicted to be important for visuomotor accuracy, we performed *in silico* cumulative ablation^49^ of all 44 morphotypes—an approach currently infeasible *in vivo* due to unavailable genetic markers. The cumulative ablation was carried out by iteratively removing one morphotype at a time and quantifying the resulting drop in accuracy while keeping previously ablated morphotypes silenced; morphotypes were ranked by the magnitude of the accuracy decrease upon removal, with the largest drops indicating the most critical contributors. For looming-evoked TPN-E response accuracy, i-TINs with dendritic and axonal arborizations restricted to the S12 layer (termed “i-S12 TIN”) and i-TINs with somata located in the OT neuropil (“SIN”) emerged as the most critical morphotypes (Figure 2F). This finding was further supported by the specific ablation of both i-S12 TINs and SINs, which resulted in a greater accuracy reduction compared with the non-specific ablation of all other TIN morphotypes (Figure 2G; *P* < 0.001, n = 15 trials). The observed accuracy decline resulted from a larger increase in looming-evoked TPN-O responses relative to TPN-E responses (Figure 2H; *P* < 0.001, n = 15 trials).

Regarding the accuracy of SMD-induced TPN-O responses, i-TINs with dendrites in the S12 layer and axons in the S56 layer (“i-S12_S56 TIN”) and i-TINs with dendrites in the S12 layer and axons in the S56 and SGC layers (“i-S12_S56_SGC TIN”) were predicted to be among the most important (Figure 2I). Consistently, the specific ablation of these TINs caused a marked reduction of accuracy compared to non-specific ablation (Figure 2J; *P* < 0.001, n = 15 trials). This reduction was attributed to a larger increase in SMD-evoked TPN-E responses relative to TPN-O responses (Figure 2K; *P* < 0.001, n = 15 trials).

These results demonstrate that specific i-TIN morphotypes ensure visuomotor accuracy by suppressing task-unrelated pathways. Specifically, i-TINs with axons terminating at relatively superficial layers suppress looming-unrelated pathways during looming processing, whereas i-TINs with axons terminating at relatively deep layers preferentially suppress SMD-unrelated pathways during SMD detection. Such targeted suppression constitutes a “push”-like mechanism that enhances visuomotor response accuracy.

### Excitatory TIN morphotypes confer visuomotor robustness primarily by enhancing task-related pathway responses

To investigate the neural substrate underlying the robustness of visuomotor transformation under noisy conditions, we first characterized the impact of escalating external noise on RGC activity. Calcium activities of RGC axonal arbors in six OT neuropil laminae (from SO to SAC) in 6-8 dpf larvae were recorded in response to looming stimuli with increasing noise levels (nl = 0, 0.01, 0.05, 0.1, 0.2, 0.5; Figure 3A). Analysis of these activities revealed a progressive decline in the signal-to-noise ratio (SNR) of RGC responses with increasing noise across all laminae, whereas S56 RGCs consistently maintained a relatively high SNR (Figure 3B). SNR was quantified as the ratio of the area under the curve (AUC) for stimulus-evoked response to that of baseline activity (see STAR Methods). When these noise-corrupted RGC activities were used to drive the model, TPN-E responses decreased with increasing noise levels, while TPN-O remained largely unresponsive (Figure 3C).

**Figure 3.**
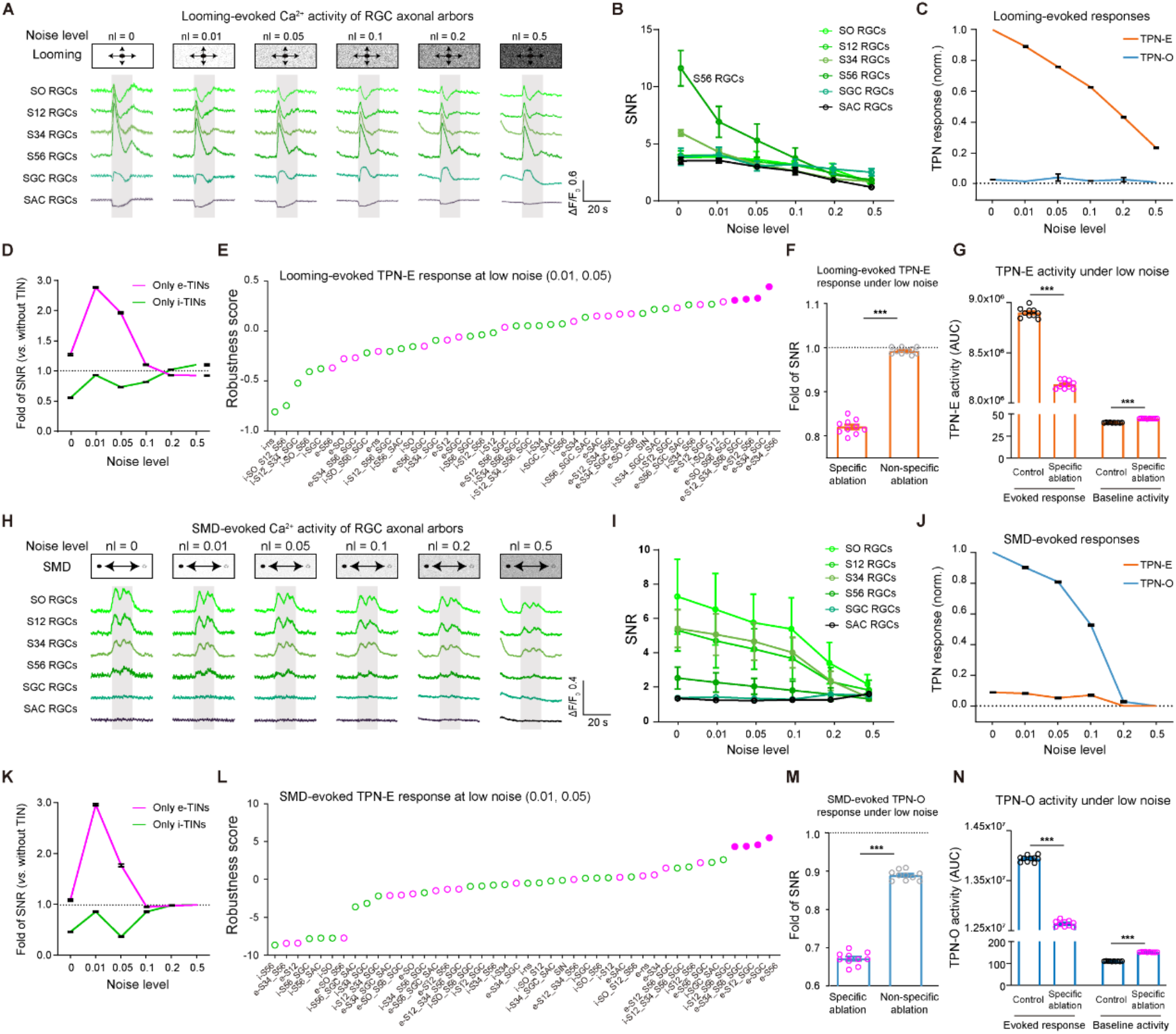
Excitatory TIN morphotypes support noise-robust visuomotor transformation. (A) Layer-specific calcium activities of RGC axonal arbors in the OT neuropil evoked by looming stimuli under increasing noise levels (nl = 0, 0.01, 0.05, 0.1, 0.2, 0.5). Top, looming stimuli; bottom, evoked calcium activities. Grey shading indicates the stimulus window. Traces represent the mean ± SEM averaged from 6 larvae (2 trials per larva); error bars are too small to be visible. (B) Signal-to-noise ratio (SNR) of layer-specific RGC responses to looming across noise levels. SNR was calculated as the ratio of evoked response AUC to that of baseline activity. Data from 6 larvae (2 trials per larva). (C) Looming-evoked TPN-E (orange) and TPN-O (blue) responses under different noise levels. The responses were normalized by the TPN-E response at nl = 0. The data were collected from 5 simulation trials for each noise level; the black bar indicates SEM. (D) Effects of e-TIN-only (magenta) and i-TIN-only (green) networks on looming-evoked TPN-E SNR across noise levels. Fold-change was computed relative to the model lacking the TIN network. The data were collected from 5 simulation trials for each noise level; the black bar indicates SEM. (E) Robustness scores of each morphotype for looming-evoked TPN-E responses under low noise levels (nl = 0.01, 0.05), revealing the top four critical e-TIN morphotypes (filled magenta dots). (F) Change in looming-evoked TPN-E SNR following ablation of the top four critical e-TIN morphotypes (Specific ablation) or ablating all other e-TIN morphotypes (Non-specific ablation). Each point denotes one simulation trial (n = 10). (G) Looming-evoked responses (left two bars) and baseline activities (right two bars) of TPN-E before specific ablation (Control black circles) or after specific ablation of the top four e-TINs (Specific ablation, magenta circles). Each point denotes one simulation trial (n = 10). (H) Layer-specific calcium activities of RGC axonal arbors in the OT neuropil evoked by SMD (4° in diameter) stimuli under increasing noise levels. Traces represent the mean ± SEM averaged from 7 larvae (2 trials per larva). (I) SNR of layer-specific RGC responses to SMD across noise levels. SNR was calculated as the ratio of evoked response AUC to that of baseline activity. Data from 7 larvae (2 trials per larva). (J) SMD-evoked TPN-E (orange) and TPN-O (blue) responses under different noise levels. The responses were normalized by the TPN-O response at nl = 0. The data were collected from 5 simulation trials for each noise level; the black bar indicates SEM. (K) Effects of e-TIN-only (magenta) and i-TIN-only (green) networks on SMD-evoked TPN-O SNR across noise levels. Fold-change was computed relative to the model lacking the TIN network. The data were collected from 5 simulation trials for each noise level; the black bar indicates SEM. (L) Robustness scores of each morphotype for SMD-evoked TPN-O responses under low noise levels (nl = 0.01, 0.05), revealing the top four critical e-TIN morphotypes (filled magenta dots). (M) Change in SMD-evoked TPN-O SNR following ablation of the top four critical e-TIN morphotypes (Specific ablation) or ablating all other e-TIN morphotypes (Non-specific ablation). Each point denotes one simulation trial (n = 10). (N) SMD-evoked responses (left two bars) and baseline activities (right two bars) of TPN-O before specific ablation (Control black circles) or after specific ablation of the top four e-TINs (Specific ablation, magenta circles). Each point denotes one simulation trial (n = 10). Each data point represents one simulation trial in F, G, M, and N. **\*\*\****P* < 0.001 (two-sided Wilcoxon signed-rank test). See also Figure S5.

We then assessed the contribution of the TIN network to this computational robustness. Comparisons with a model lacking the TIN network revealed that the inclusion of all e-TIN morphotypes enhanced the SNR of TPN-E responses under low-noise conditions (nl = 0.01, 0.05), whereas the incorporation of all i-TIN morphotypes reduced TPN-E SNR to a lesser extent (Figure 3D). To pinpoint the specific TIN morphotypes responsible for this external noise resilience, we assessed the contribution of each TIN morphotype by enabling only that morphotype within an otherwise silent TIN layer and quantified its effect on TPN-E SNR. The impact of each morphotype on TPN-E SNR was quantified as fold-change relative to the model lacking the TIN network under no or low-noise conditions (Figures S5A-S5C). To specifically evaluate their capacity to counteract external noise, a noise-robustness score was computed by subtracting the fold-change at nl = 0 from the average fold-change at nl = 0.01 and 0.05. This analysis identified a subset of e-TIN morphotypes with axons terminating at deep neuropil layers as the primary contributors to noise robustness (Figure 3E).

The model-inferred necessity of these top-contributing morphotypes was then tested by specifically ablating the four highest-ranked morphotypes within a model incorporating the TIN network. This selective ablation induced a significantly larger reduction in TPN-E SNR compared with the ablation of the remaining 16 e-TIN morphotypes (Figure 3F; *P* < 0.001), supporting a model-predicted role of this specific group in maintaining visuomotor robustness against external noise. Further analysis showed that the dominant effect was a reduction in the looming-evoked response, accompanied by a smaller increase in baseline activity (Figure 3G; *P* < 0.001).

A parallel set of experiments using SMD stimuli, which drive the TPN-O orienting pathway, revealed a similar computational strategy for ensuring visuomotor robustness. In response to SMDs with increasing noise levels, RGC arborizations in superficial layers (particularly SO) exhibited a high SNR compared to other layers (Figures 3H and 3I). Modeling showed that TPN-O responses markedly decreased with rising noise, while TPN-E responses remained minimal (Figure 3J). As with the TPN-E escape pathway, e-TINs provided greater benefit to the SNR of TPN-O responses than i-TINs (Figure 3K). A similar morphotype-assessing procedure identified a distinct ensemble of e-TINs with axons arborizing at deep layers most critical (Figures 3L and S5D-S5F). Ablation of the top-contributing group (4 morphotypes) resulted in a more pronounced reduction in TPN-O SNR compared to non-specific ablation (Figure 3M; *P* < 0.001). This reduction was attributable to a substantial decrease in SMD-evoked responses alongside an increase in baseline activity (Figure 3N; *P* < 0.001).

These results demonstrate that specific e-TIN morphotypes confer visuomotor robustness by amplifying task-related pathway responses. For each behaviorally distinct circuit—looming-evoked escape and SMD-induced orienting—a distinct ensemble of e-TIN morphotypes with axons terminating at deep layers is engaged. Such targeted enhancement implements a “pull”-like mechanism that stabilizes visuomotor response against external noise, showcasing a fundamental neural algorithm for reliable computation within biological networks.

### OT-projecting pretectal 5-HT neurons confer visuomotor flexibility by reweighting competing pathways

Neural systems face a fundamental stability-flexibility trade-off. Although long-term plasticity mechanisms enable behavioral adaptation, their metabolic cost and temporal inertia constrain real-time flexibility.^50–52^ Neuromodulatory systems offer a solution by rapid, energy-efficient circuit reconfiguration.^53–55^ Among these, phylogenetically conserved 5-HT systems play a crucial role in modulating sensorimotor processing,^56–59^ with axonal projection to the OT observed across vertebrates.^60,61^

To examine the role of 5-HT systems in visuomotor flexibility, we first mapped the single-cell morphology of 5-HT neurons in 6 - 8 dpf larval zebrafish (see STAR Methods), and reconstructed 36 5-HT neurons with somata located in the pretectum and axonal projections innervating the OT neuropil (Figure 4A, left). Their collective axons preferentially target superficial and deep neuropil layers (Figure 4A, right), which overlap with the superficial and deep OT domains preferentially engaged by SMD- and looming-related inputs, respectively.^17^ Based on axonal projection patterns, we classified them into two subtypes: deep layers-projecting 5-HT (DL_5-HT) neurons preferentially targeting SGC and S56 layers (Figure 4B), and superficial layers-projecting 5-HT (SL_5-HT) neurons primarily arborizing in SO and S12 layers (Figure 4C). Given that 5-HT receptors are commonly expressed on both pre- and post-synaptic sites,^62^ we quantified putative connectivity between these pretectal 5-HT neurons’ axons and the dendritic/axonal compartments of TNs (including TINs and TPNs) based on close appositions (see STAR Methods). DL_5-HT and SL_5-HT neurons showed differential connectivity patterns with TINs, and both subtypes tended to exhibit relatively higher connection probabilities with TPNs than with TINs (Figure 4D and Table S3). We then examined the potential effect of 5-HT neurons on visuomotor transformation by performing *in vivo* cell-attached recording. Bath application of 5-HT increased the spontaneous firing of TNs (Figure 4E; *P* < 0.01), suggesting that 5-HT neurons enhance TN excitability. This regulatory effect was then incorporated into the SNN model based on the connectivity pattern between 5-HT neurons and TNs.

**Figure 4.**
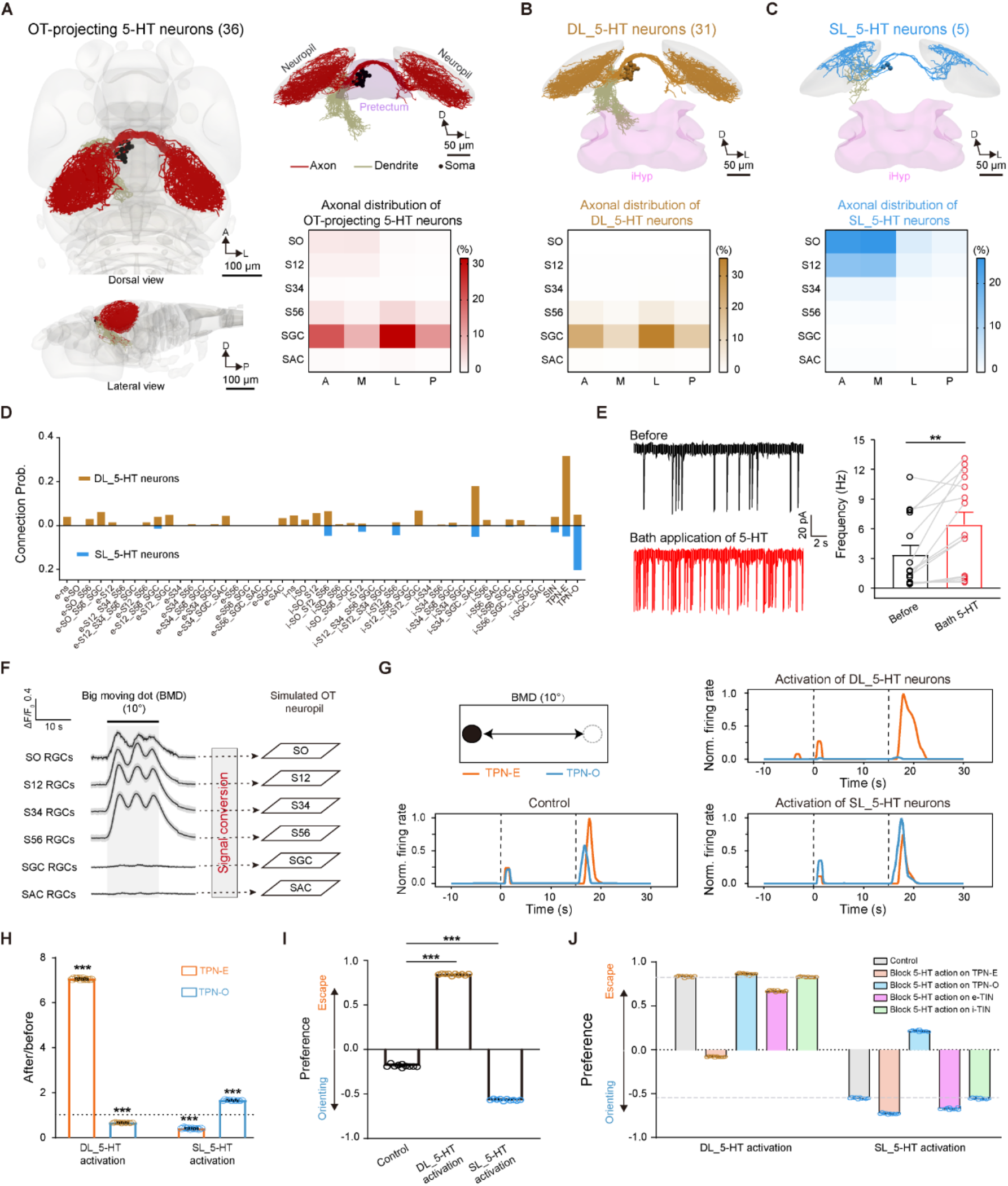
OT-projecting pretectal 5-HT neurons enable flexible bias of visuomotor outputs. (A) Reconstructed single-cell morphologies of OT-projecting pretectal 5-HT neurons. Left, dorsal and lateral views of OT-projecting 5-HT neurons. Right, dorsal-posterior view of 5-HT neurons (top) and normalized distribution of their axonal arbors within the OT neuropil (bottom). Normalized axonal distribution was calculated by dividing the axonal projection length in each microdomain by total axonal projection length in the OT neuropil. Numbers in parentheses indicate the number of reconstructed 5-HT neurons. All 5-HT neurons were mirrored to the left hemisphere for visualization. (B) Pretectal 5-HT neurons projecting to deep layers of the OT neuropil (DL_5-HT neurons). Top, dorsal-posterior view of DL_5-HT neurons. Bottom, normalized axonal distribution of DL_5-HT neurons in the OT neuropil. The number in parentheses indicates the number of reconstructed DL_5-HT neurons. Some dendrites arborize in the intermediate hypothalamus (iHyp). (C) Pretectal 5-HT neurons projecting to superficial layers of the OT neuropil (SL_5-HT neurons). Top, dorsal-posterior view of SL_5-HT neurons. Bottom, normalized axonal distribution of SL_5-HT neurons in the OT neuropil. The number in parentheses indicates the number of reconstructed SL_5-HT neurons. (D) Putative connection probability of DL_5-HT and SL_5-HT neuronal axons to TINs and TPNs (see Table S3). (E) Effect of bath application of 10 μM 5-HT on TN activity. Left, typical *in vivo* whole-cell recording of TNs before and during 5-HT application. Right, summary of spontaneous firing frequency. Each dot represents one recorded trial. Repeated trials were averaged within neuron before statistical testing (n = 3 neurons, 5 trials per neuron). **\*\****P* < 0.01 (two-sided Wilcoxon signed-rank test). (F) Layer-specific calcium activities of RGC axonal arbors in the OT neuropil evoked by big moving dot (BMD, 10° in diameter). These calcium activities served as inputs to the OT SNN model after conversion into spike-proxy current inputs. Left, BMD-evoked RGC responses. Right, schematic showing that RGC activity is fed into the model. Activity traces represent the mean ± SEM averaged from 7 larvae (2 trials per larva); error bars are too small to be visible. The imaging was performed on double-transgenic *Tg(-7atoh7:GAL4-VP16);Tg(UAS:GCaMP6s)* larvae at 6-8 dpf. (G) BMD-evoked TPN-E and TPN-O responses under control conditions without 5-HT neuronal activation (bottom left), with activation of DL_5-HT neurons (top right), or with activation of SL_5-HT neurons (bottom right). Top left, schematic of the BMD stimulus, which lasts 15 s. The window between the two dashed lines indicates the visual stimulation period. Traces are averaged from 5 simulation trials. (H) DL_5-HT and SL_5-HT neuronal activation-induced changes in BMD-evoked TPN-E and TPN-O responses. Each point denotes one matched simulation trial (n = 10). The dashed line indicates the no-change level (fold-change = 1). **\*\*\****P* < 0.001 (two-sided one-sample Wilcoxon signed-rank tests against 1 for each group). (I) DL_5-HT and SL_5-HT neuronal activation-induced changes in BMD-evoked behavioral preference. The preference index was calculated as (AUC of TPN-E minus AUC of TPN-O) divided by the sum of both AUCs. n = 10 matched simulation trials. (J) Component-wise blocking analysis of behavioral preference shifts induced by DL_5-HT or SL_5-HT neuronal activation, revealing differential contributions of OT neuron classes. A, anterior; L, lateral; D, dorsal; P, posterior.

We then investigated whether the activation of these pretectal 5-HT neurons can bias behavioral outputs under ambiguous sensory conditions. We presented the model with RGC activities evoked by a large black moving dot (BMD, 10° in diameter; Figure 4F), a neutral stimulus that does not consistently evoke escape or orienting responses.^41,63^ In the model, BMD activated both TPN-E and TPN-O (Figure 4G, bottom left). However, activating DL_5-HT neurons significantly enhanced TPN-E responses but suppressed TPN-O activity (Figures 4G, top right and 4H, left two bars; *P* < 0.001). Conversely, activation of SL_5-HT neurons largely enhanced TPN-O responses while suppressing TPN-E activity (Figures 4G, bottom right and 4H, right two bars; *P* < 0.001). Consequently, compared to the control group, DL_5-HT neuron activation induced a stronger escape preference, while SL_5-HT neuron activation caused a stronger orienting preference (Figure 4I; *P* < 0.001). Here, the behavioral preference index was computed as the difference between TPN-E and TPN-O AUCs divided by their sum. Similar pathway-biased modulation was observed for a larger ambiguous stimulus (16° BMD). Under control conditions, the model showed co-activation of TPN-E and TPN-O, whereas activation of DL_5-HT neurons preferentially enhanced TPN-E responses to drive an escape bias, and activation of SL_5-HT neurons boosted TPN-O responses to induce an orienting bias (Figures S6A-S6D). These results indicate that this pretectal 5-HT system confers flexible bias over visuomotor outputs.

To identify which neuronal types within the OT mediate this flexible modulation, we blocked *in silico* the effect of 5-HT neuron activation on specific TN classes, including TPN-E, TPN-O, i-TIN, and e-TIN morphotypes. The model showed that the escape preference induced by DL_5-HT neuron activation was primarily mediated by TPN-E, while the orienting preference induced by SL_5-HT neuron activation was mainly mediated by TPN-O, with TINs making minor contributions in both conditions (Figures 4J and S6E). These simulations suggest that the flexible bias is mediated mainly by 5-HT-dependent gain changes in TPN-E and TPN-O, with smaller contributions from TINs.

### Biological validation of the roles of OT-projecting 5-HT subsystems in visuomotor transformation

To examine whether such flexible modulation occurs in animals, we first conducted *in vivo* calcium imaging of the axonal arbors of DL_5-HT and SL_5-HT neurons within the OT neuropil and found they displayed distinct response patterns (Figures 5A and 5B). While SL_5-HT neurons were tonically activated during sustained background illumination, DL_5-HT neurons displayed transient activity in response to both light onset and offset (Figure 5B). When faced with BMD stimulus, two populations displayed antiphasic activity patterns (Figure 5C). Consistent with this pattern, simulations revealed that tuning the relative activity levels between DL_5-HT and SL_5-HT neurons progressively reweighted the TPN-E and TPN-O pathways, thereby driving a graded shift in behavioral preference (Figure 5D).

**Figure 5.**
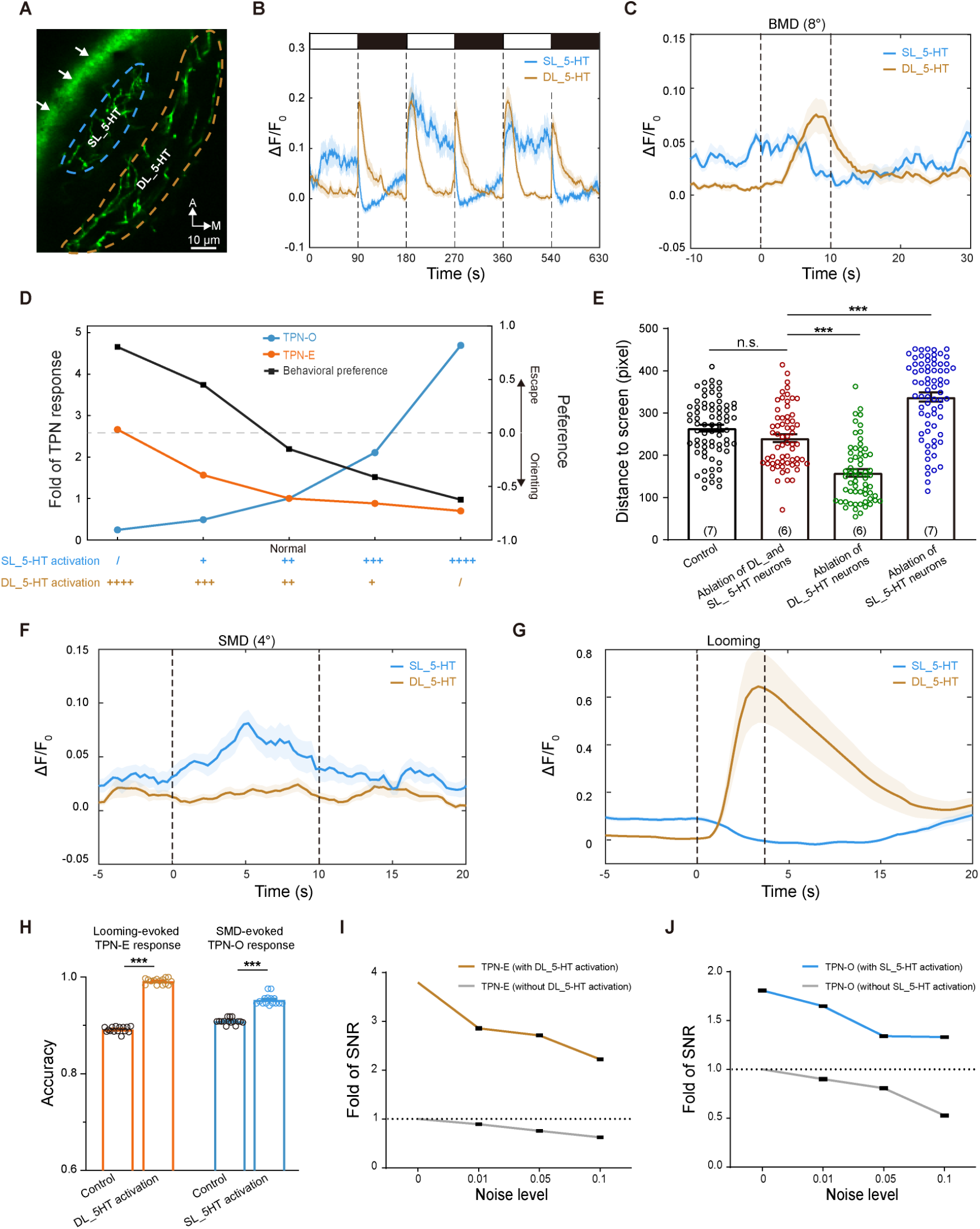
Pretectal 5-HT subsystems bias adaptive visuomotor behavior and increase accuracy/robustness under related visual stimuli. (A) Example imaging field showing the region-of-interest (ROI) masks used to separately quantify DL_5-HT and SL_5-HT axonal signals within the OT neuropil. The white arrows indicate non-signaling fluorescence on the skin. (B) Calcium responses of DL_5-HT and SL_5-HT axonal signals during repeated light-dark transitions. White and black bars above the traces indicate light and dark periods, respectively. Activity traces represent mean ± SEM across 7 larvae. (C) Calcium responses of DL_5-HT and SL_5-HT axonal signals to BMD (8° in diameter) stimulation. The window between the two dashed lines indicates the visual stimulation period. Traces represent mean ± SEM across 7 larvae. (D) Simulated fold changes in TPN-E response (orange), TPN-O response (blue), and behavioral preference (black) across a graded shift of the activity dominance from DL_5-HT to SL_5-HT neurons. The “Normal” denotes the activity level of DL_5-HT and SL_5-HT neurons in response to BMD (8° in diameter). Positive preference values indicate escape bias, whereas negative values indicate orienting bias. (E) Effect of *in vivo* ablation of DL_5-HT, SL_5-HT, or both DL_5-HT and SL_5-HT neurons on BMD-evoked behaviors in 7-dpf larvae. Distance was measured as the distance from the fish head to the screen presenting BMD stimuli. A smaller distance indicates that the fish approached the screen, implying orienting behavior, while a larger distance indicates that the fish moved away from the screen, implying escape behavior. Each dot represents one behavioral trial. For statistical testing repeated trials from the same larva were averaged to obtain one value per larva. Bars show the mean and error bars indicate SEM; numbers on the bars indicate the number of larvae examined. n.s., no significance; **\*\*\****P* < 0.001 (two-sided Mann-Whitney U tests). (F) Calcium responses of DL_5-HT and SL_5-HT axonal signals to SMD (4° in diameter) stimulation. The window between the two dashed lines indicates the visual stimulation period. Traces represent mean ± SEM across 7 larvae. (G) Calcium responses of DL_5-HT and SL_5-HT axonal signals to looming stimulation. The window between the two dashed lines indicates the visual stimulation period. Traces represent mean ± SEM across 5 larvae. (H) Changes in looming-evoked TPN-E response accuracy and SMD-evoked TPN-O response accuracy induced by DL_5-HT or SL_5-HT activation. Each circle denotes one simulation trial (n = 15). (I) Fold change in the SNR of looming-evoked TPN-E responses relative to the control group across increasing noise levels, with or without DL_5-HT neuronal activation. The “without DL_5-HT activation” trace corresponds to the control TPN-E response shown in Figure 3C. DL_5-HT neuronal activation increases the SNR of the escape pathway across noise levels. The data were collected from 5 simulation trials for each noise level; the black bar indicates SEM. (J) Fold change in the SNR of SMD-evoked TPN-O responses relative to the control group across increasing noise levels, with or without SL_5-HT neuronal activation. The “without SL_5-HT activation” trace corresponds to the control TPN-O response shown in Figure 3J. SL_5-HT neuronal activation increases the SNR of the orienting pathway across noise levels. The data were collected from 5 simulation trials for each noise level; the black bar indicates SEM.

We then selectively ablated bilateral DL_5-HT, SL_5-HT neurons, or both populations in larval zebrafish based on their distinct visual response patterns (see Methods). Compared to control and both population ablation groups, ablation of only DL_5-HT neurons caused larvae to prefer approaching the screen presenting BMD, whereas ablation of only SL_5-HT neurons caused larvae to prefer escaping away from it (Figure 5E; see STAR Methods). These behavioral results are consistent with the model predictions, supporting a mechanism in which OT lamina-specific pretectal 5-HT subsystems enable visuomotor flexibility by reweighting competing TPN-E and TPN-O pathways.

Given that the 5-HT subsystems exhibited ecologically relevant, context-dependent activity patterns, with SL_5-HT and DL_5-HT neurons responding preferentially to prey-like SMDs and predator-mimicking looming stimuli (Figures 5F and 5G), respectively, we tested whether activating the corresponding 5-HT subsystem could further bias its target pathway. Simulations showed that DL_5-HT neuron activation improved looming-evoked TPN-E accuracy and robustness (Figures 5H, left and 5I), whereas SL_5-HT neuron activation elevated SMD-evoked TPN-O accuracy and robustness (Figures 5H, right and 5J), suggesting that the OT-projecting 5-HT subsystems additionally contribute to visuomotor accuracy and robustness via pathway-specific gain modulation.

## DISCUSSION

By combining *in silico* perturbations with biological experiments, we provide evidence that the OT implements adaptive visuomotor transformation through the complementarity of intrinsic interneuron circuits and extrinsic neuromodulatory systems. First, we identified a morphotype-resolved tectal “push-pull”-like architecture, in which i-TINs suppress task-unrelated pathways while e-TINs reinforce task-related pathways. In response to ecologically relevant stimuli (Figure 6A), visuomotor accuracy is promoted by i-TINs that preferentially suppress responses in the task-unrelated pathway. Concurrently, visuomotor robustness is supported by e-TINs that amplify responses of the task-related pathway. Second, under ambiguous stimuli (Figure 6B), visuomotor flexibility is enabled by visually responsive, lamina-specific 5-HT subsystems that bias competition between visuomotor pathways by modulating tectal output neurons (i.e., TPNs).

**Figure 6.**
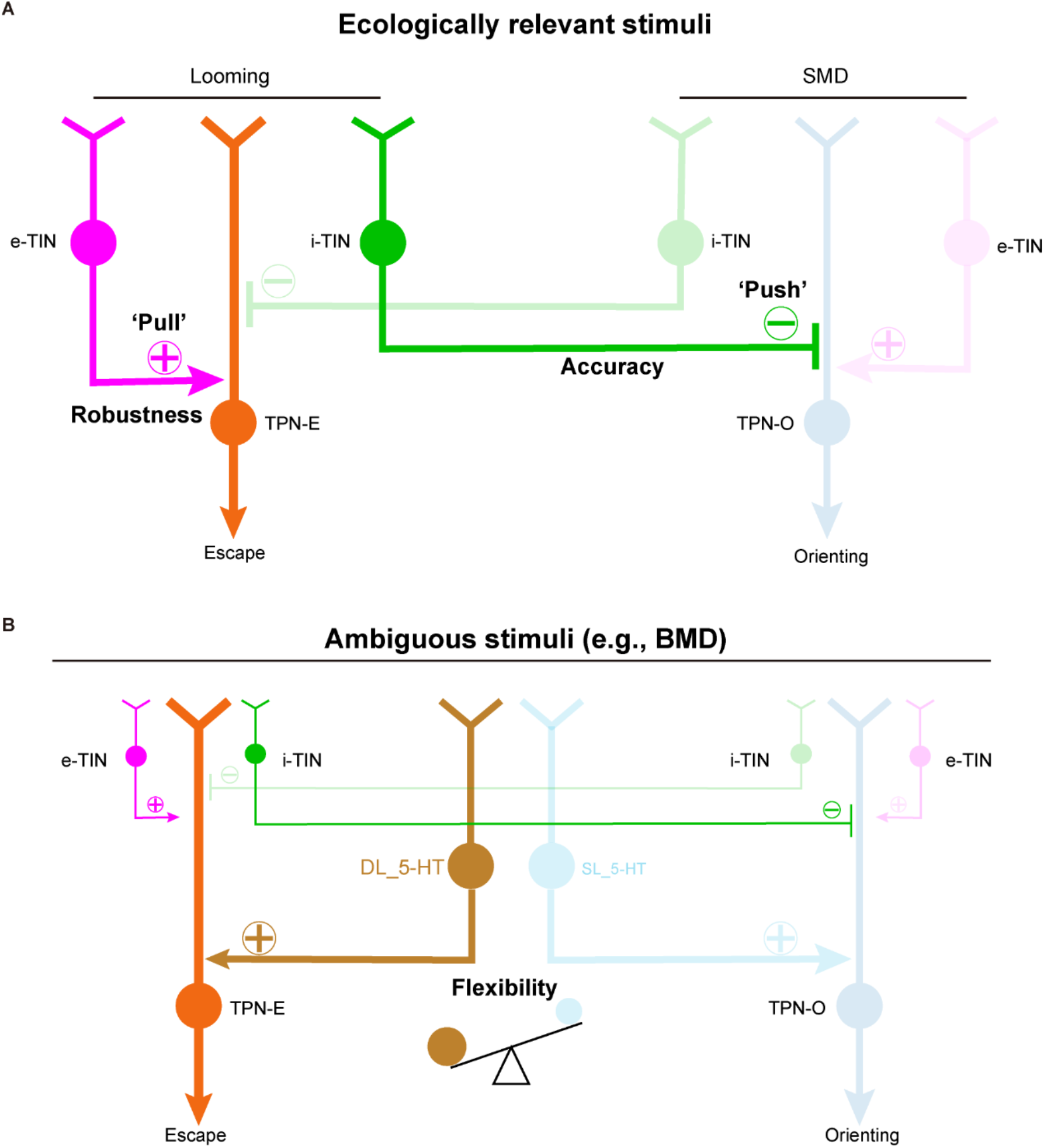
Schematic model of adaptive visuomotor transformation. (A) In ethologically relevant contexts, i-TINs improve visuomotor accuracy by suppressing task-unrelated pathways (“Push”), whereas e-TINs improve visuomotor robustness by amplifying task-relevant pathway responses (“Pull”). (B) Under ambiguous contexts, lamina-specific serotonergic subsystems bias pathway selection by modulating TPN activity: DL_5-HT biases the circuit toward escape output (TPN-E), whereas SL_5-HT biases it toward orienting output (TPN-O). Balanced engagement of these subsystems enables flexible behavioral selection.

Beyond the identified push-pull motif, our results support the TIN network functioning as a biologically structured reservoir. In this context, a reservoir refers to a fixed, sparse, heterogeneous recurrent circuit capable of transforming distributed retinal inputs into pathway-selective outputs while sustaining robust computation under noisy sensory drives. Structurally, the tectum embodies these features: its recurrent TIN core compose of 44 morphotypes, whose connectivity is sparse, inhibitory-dominant, and topologically distinct from random baselines (see Figures 2C and S4H-S4N). Functionally, the recurrent core is necessary for the pathway selectivity, since feedforward bias alone was insufficient to explain the selective responses observed with the TIN layer intact, but abolished upon its removal (see Figures 1I, 1J, and 2C-2E). At dynamical level, the TIN network displays reservoir-like dynamics, specifically stable perturbation decay and finite-delay fading memory (see Figures S4O and S4P). In addition, this substrate provides morphotype-specific support for response accuracy (see Figure 2) and noise robustness (see Figure 3). These lines of evidence support that the TIN circuit functions as a biological substrate for reservoir computing during visuomotor transformations.

The diversity of interneurons underlies the computational capacity and functional versatility of neural circuits.^5^ But the limited genetic access has hindered causal dissection of most interneuron subtypes. In the present study, we integrated morphotype-resolved *in silico* perturbations and showed that morphologically defined interneuron types serve as push-pull-like circuit motifs for visuomotor transformation. For accuracy, i-TINs provide layer-matched suppression of task-unrelated pathways: the laminar depth of i-TIN axonal arbors aligns with the retinorecipient locations where unrelated RGC inputs terminate (comparing the distributions of stimulus-evoked responses and involved i-TIN’ axon arbors in Figures 2B, 2F, and 2I), thereby selectively attenuating task-unrelated pathways while preserving the related drive. This lamina-specific “push” can sharpen pathway selection and improve visuomotor accuracy. In contrast, robustness relies predominantly on e-TINs with axons arborizing in deep layers, where they connect with TPN-E and TPN-O (see Table S2), providing a stable recurrent “pull” that reinforces the task-related pathway under degraded sensory conditions. Thus the laminar complementarity between i-TIN-mediated suppression and e-TIN-mediated reinforcement offers a circuit-level account for how the tectal reservoir achieves both accurate and robust transformation via push-pull-like mechanisms. Recent advances in targeting diverse OT interneuron morphotypes can provide opportunities to test these model-derived predictions *in vivo* in the future.^45^

Neuromodulatory systems can rapidly reconfigure circuit function without requiring structural rewiring.^64^ We found that lamina-specific 5-HT neurons can bias visuomotor outputs even when OT connectivity remains unchanged: a deep-layer-projecting 5-HT subsystem biases the circuit toward escape, whereas a superficial-layer-projecting subsystem biases it toward orienting. Under natural contexts, both 5-HT subsystems can be antiphasically activated by neutral stimuli (e.g., BMD) to varying degrees, dynamically reweighting escape and orienting readouts and shifting behavioral preference. This lamina-specific gain modulation echoes broader evidence that serotonergic systems can bias action selection and behavioral persistence across contexts and is conceptually consistent with recent recurrent-network studies of neuromodulatory control.^58,65,66^

In summary, our findings elucidate how cellular-level mechanisms, spanning interneuron morphotypes to neuromodulatory systems, enable the OT to transform visual inputs into accurate, robust, and flexible motor-related outputs. These insights advance our understanding of vertebrate visuomotor transformation and suggest design principles for brain-inspired computational systems that operate robustly in noisy and dynamic environments.

### Limitation of the study

This study has several limitations. First, the neural connections used in the model were inferred from close appositions within the single-cell morphology-based zebrafish atlas,^18^ as brain-wide synapse-resolved complete reconstructions are not yet available, despite the emergence of larval zebrafish electron microscopy resources.^67–73^ Second, the reservoir interpretation is based on mesoscopic connectivity, model-based reservoir-state analyses, and task-specific behavior rather than direct *in vivo* measurement of the full TIN population dynamics across all possible regimes; future work should test perturbation decay, temporal memory, and readout geometry directly with population recordings and targeted morphotype perturbations. Third, the spiking model employs simplified neuron and synapse dynamics. Fourth, serotonergic modulation was implemented as a net increase in neuronal excitability, whereas 5-HT can act through multiple receptor types with compartment- and cell-type-specific effects. Future work integrating synapse-resolved connectomes, morphologically detailed compartmentalized modeling, and 5-HT receptor-specific perturbations as well as targeted manipulation of identified morphotypes will be essential to test and refine the proposed push-pull-like and neuromodulatory bias mechanisms.

## RESOURCE AVAILABILITY

### Lead contact

Further information and requests for resources and reagents should be directed to and will be fulfilled by the lead contact, Jiu-Lin Du.

### Materials availability

Zebrafish lines and plasmids generated or maintained by the Jiu-Lin Du lab are available from the lead contact upon reasonable request, subject to institutional and material transfer agreements.

### Data and code availability

All data generated or analyzed during this study are included in this manuscript and its supplemental information. Source data for the figures are provided with this paper. All mesoscopic connectome data used in this study, including the raw files for optic tectum neurons, serotonergic (5-HT) neurons, and retinal ganglion cells (RGCs), are available from the CEBSIT Brain Science Data Archive at https://www.braindatacenter.cn/datacenter/web/index.html#/dataSet/details?id=1585563873913057281. The foundational zebrafish mesoscopic atlas data^18^ used in this study are also publicly accessible through the ZExplorer web portal at https://zebrafish.cn/LM-Atlas/EI. All other data are available from the corresponding authors upon request. The custom code and analysis scripts used to construct the spiking neural network model, perform *in silico* ablations, and analyze network dynamics are publicly available at https://github.com/yqian-CEBSIT/Optic_Tectum_Simulation and are released under the MIT License. The BrainPy framework^74^ used for simulations is publicly available at https://github.com/brainpy/BrainPy.

## ACKNOWLEDGMENTS

We thank Drs. Chao-Ming Wang, Tian-Qiu Zhang, Si Wu and Yu Mu for suggestions and support. We appreciate the technical assistance from Drs. Hui Zhang and Qian Hu. We acknowledge the assistance with neuronal morphology registration on the brain template from the Facility of Mapping Zebrafish Brain-Wide Mesoscale Connectome at CEBSIT for technical support. This work was supported by Brain Science and Brain-like Intelligence Technology - National Science and Technology Major Project (2021ZD0204500, 2021ZD0204502, and 2022ZD0209600), National Natural Science Foundation of China (32321003, and U22B2063), Shanghai Municipal Science and Technology Projects (25511102500 and 2025SHZDZX025D03), Shanghai Municipal Commission of Economy and Informatization (2025-GZL-RGZN-BTBX-02005), and Key Research Program of Frontier Sciences (QYZDYSSW-SMC028) and Strategic Priority Research Program (XDB32010200) of Chinese Academy of Sciences.

## AUTHOR CONTRIBUTIONS

Conceptualization: Y.Q., and J.D.; Code writing and brain simulation: Y.Q.; Methodology: Y.Q.; Investigation: Y.Q., S.L., and J.D.; Data acquisition and pre-processing of wet experiments (including neuron morphology imaging and reconstruction, calcium imaging, and electrophysiology): S.L., Y.Q., M.C., H.W., T.Z. and X.D.; Funding acquisition: J.D. and X.D.; Project administration: Y.Q., S.L., and J.D.; Supervision: J.D., and X.D.; Manuscript writing: Y.Q., S.L., and J.D. with inputs from others.

## SUPPLEMENTAL INFORMATION

Supplemental information includes Figures S1-S6 and Tables S1-S3.

## STAR METHODS

### KEY RESOURCES TABLE

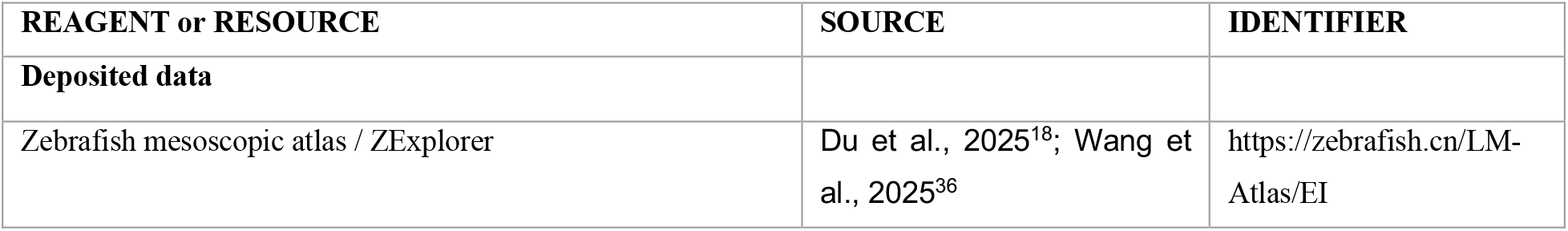
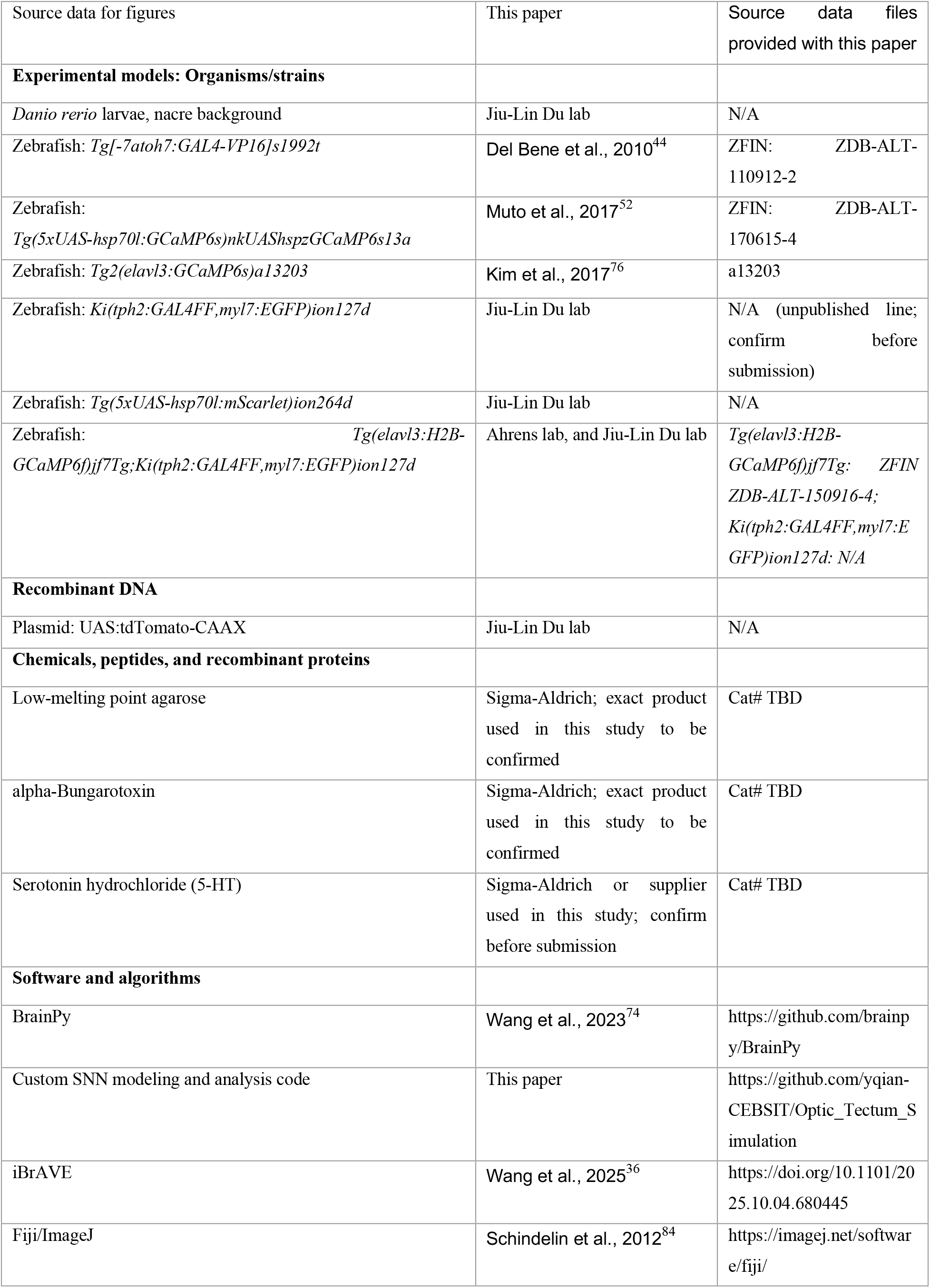
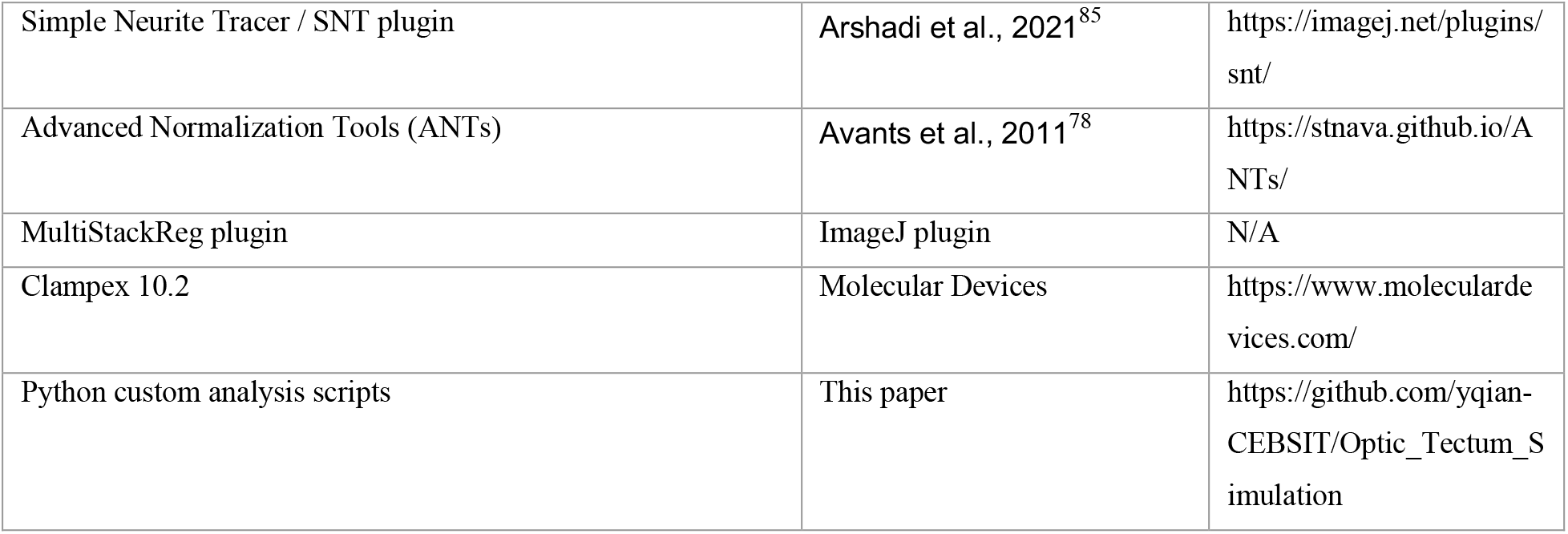

### EXPERIMENTAL MODEL AND STUDY PARTICIPANT DETAILS

#### Zebrafish husbandry and preparation

Zebrafish larvae were maintained under standard conditions at 28.5°C with a 14-h light/10-h dark photoperiod. Larvae were kept in Hank’s solution until 2 dpf, and then transferred to facility water. Beginning at 5 dpf, larvae were fed rotifers once daily. All experiments were conducted using larvae aged 6 - 8 dpf.

For *in vivo* imaging experiments, larvae were embedded in 2.0% low-melting point agarose (Sigma) dissolved in facility water. After 5-min embedding, facility water was added to the recording chamber. For *in vivo* electrophysiological recordings, facility water was replaced with an external solution containing (in mM): 134 NaCl, 2.9 KCl, 2.1 CaCl_2_·2H_2_O, 1.2 MgCl_2_·6H_2_O, 10 HEPES, and 10 D-glucose (pH 7.8, 290 mOsm). All animal protocols were reviewed and approved by the Animal Care and Use Committee of the Center for Excellence in Brain Science and Intelligence Technology, Chinese Academy of Sciences (NA-046-2023).

#### Zebrafish transgenic lines

Transgenic zebrafish larvae were in a nacre background.^75^ Fishline *Tg(-7atoh7:GAL4-VP16)s1992t* (in abbreviation: *Tg(-7atoh7:GAL4-VP16*)),^44^ *Tg(UAS:GCaMP6s)nkUAShspzGCaMP6s13a* (*Tg(UAS:GCaMP6s)*),^52^ *Tg2(elavl3:GCaMP6s)a13203* (*Tg2(elavl3:GCaMP6s)*),^76^ *Ki(tph2:GAL4FF,myl7:EGFP)ion127d* (*Ki(tph2:GAL4FF,myl7:EGFP)*)^77^ were described in previous studies, respectively, and *Tg(5xUAS-hsp70l:mScarlet)ion264d* was constructed in the lab.

### METHOD DETAILS

#### 5-HT neuron reconstruction

We microinjected UAS:tdTomato CAAX plasmid into one cell stage fertilized egg of *Tg(HuC:H2B-GCaMP6f);Ki(tph2:GAL4FF,myl7:EGFP)* to sparsely label 5-HT neurons.^18^ 5-HT neurons were semi-automatically traced using the Simple Neurite Tracer (SNT) plugin in Fiji. The resulting neuron skeletons, composed of interconnected nodes, were saved as individual SWC files. The neuron skeleton was then registered to the common physical space^18^ using the antsApplyTransformToPoints function in ANTs.^78^

#### TIN axon and dendrite discrimination

To identify the dendritic and axonal compartments of TINs, we applied a machine-learning-based classifier trained on neuron reconstructions with dendrite-axon annotations derived from morphology and presynaptic-site labeling data.^18,36^ A subset of TINs was sparsely labeled with the presynaptic marker Synaptobrevin-GFP (Sypb-EGFP) for validation (see Figures S4A-S4C). In these neurons, discrete Sypb-EGFP puncta were localized to axonal branches, providing a direct molecular signature of the axonal compartment. The dendritic compartment was correspondingly defined as Sypb-negative processes. This ground-truth dataset allowed us to calibrate and validate the morphological criteria described in the Results.

This combined approach was systematically applied to define the polarity of all 1,186 TINs, which were classified into three canonical arborization patterns for network construction: 1) Monostratified TINs: both dendrites and axons were confined to a single, identical lamina. The axon was identified as the compartment within that lamina exhibiting finer caliber and high branching complexity; 2) Multilayered TINs: dendrites predominantly occupied superficial laminae (e.g., SO, SFGS), while axons arborized in deeper laminae (e.g., SGC, SAC). The compartment in the deeper layer, characterized by finer processes, was classified as the axon; 3) Non-stratified TINs: dendritic and axonal processes intermingled across laminae without clear spatial segregation. In these cases, the identification relied largely on the fine, complex arborization pattern of the axon.

#### Quantification of neuronal connectivity

Because spatial proximity between axonal and dendritic compartments can provide a useful proxy for synaptic connectivity across systems, we inferred putative anatomical connectivity from axon-dendrite proximity in the registered common space.^79–83^ Putative anatomical connectivity throughout this study (RGC to TPN and TIN, TPN to M-cell and nMLF, TIN to TIN, and TIN to TPN) was inferred from axon-dendrite proximity in the registered common coordinate space using the Neuronal Wiring Calculation filter in the iBrAVE platform.^36^ For each presynaptic neuron i, axonal compartments were compared to dendritic compartments of each postsynaptic neuron j, and a putative contact was assigned when the Euclidean distance (*d*) between their centroids was less than or equal to a distance cutoff. A putative connection between a neuron pair (*i, j*) was defined if at least one contact existed. Connection probability between two populations *A*(presynaptic) and *B* (postsynaptic) was then computed as:

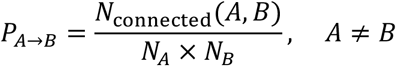

Here, *N*_connected_(*A, B*) is the number of connected ordered neuron pairs, and *N*_*A*_ and *N*_*B*_ are the numbers of neurons in the two populations. For within-population connectivity, the denominator was *N*_*A*_ × (*N*_*A*_ − 1), thereby excluding autapses.

To assess the robustness of distance-threshold selection, we generated morphotype-level connection probability matrices at *d* = 0.5, 1, 2, 4 and 6 μm. Mean pairwise similarity (Pearson’s r) across matrices peaked at 2 and 4 μm (see Figures S4E and S4F), suggesting that these cutoffs provide a stable and representative estimate of the underlying connectivity.

Matrix density was defined as the fraction of morphotype-to-morphotype entries with non-zero connection probability.

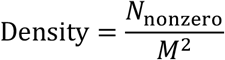

Here, *M* is the number of morphotypes in the matrix, and same-type entries on the diagonal were retained because they represent within-morphotype connectivity between distinct neurons rather than single-neuron self-connections. Density is therefore dimensionless and has a theoretical maximum of 1 when all *M*^2^ entries are non-zero.

To quantify how selectively each source morphotype distributes its outputs across target morphotypes, we computed row selectivity from the morphotype-level connection probability matrix *W* = [*w*_*ij*_]. For each source morphotype *i*, outgoing probabilities were normalized across targets, row entropy was calculated with the convention 0log0 = 0, and the entropy was then normalized by the total number of morphotypes. Row selectivity was defined as follows.

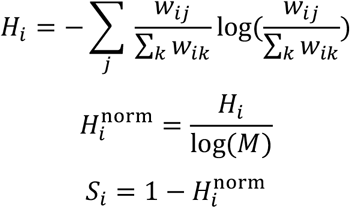

Mean row selectivity was obtained by averaging *S*_*i*_ across source morphotypes with non-zero total outgoing probability. Higher selectivity indicates that a source morphotype preferentially targets a smaller subset of postsynaptic morphotypes, whereas lower selectivity indicates more diffuse connectivity. 2 μm was selected as the cutoff throughout the study, because smaller values produced sparser matrices at the expense of false negatives, larger values increased density and reduced selectivity at the cost of false positives (see Figure S4G), and matrix similarity stabilized by this threshold.

To ensure directionality and biological plausibility, we exclusively considered axon→dendrite relationships, using axon/dendrite polarity annotations for TINs as described above. These connectome-derived probabilities were used to instantiate directed connectivity matrices for SNN construction.

#### SNN model construction

We constructed a biologically plausible SNN using BrainPy, a flexible and high-performance simulation framework for spiking dynamics and neural computation.^74^ Neuronal dynamics followed LIF formalism with absolute refractory periods. Differential equations were solved using the Euler method (dt = 0.1 ms). Parameter sets for each neuron class are in Table S1, derived from *in vivo* whole-cell recordings of TNs.

The subthreshold dynamics of membrane potential *V*(*t*) are governed by:

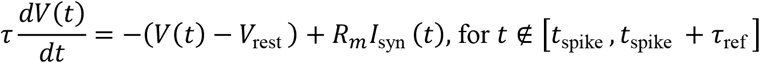

Where *V*(*t*) is the membrane potential, *V*_rest_ is the resting potential, *V*_reset_ is the reset potential after a spike, *V*_th_ is the spike threshold, τ is the membrane time constant, *τ*_ref_ is the refractory period, *R*_1_ is the membrane resistance, and *I*_syn_ (*t*) is the total synaptic current input. When *V*(*t*) ≥ *V*_th_, the neuron emits a spike and resets to *V*_reset_, remaining inactive for the duration of *τ*_ref_. The detailed parameterization for each modeled neuron type is listed in Table S1.

For synaptic model, synaptic transmission was implemented as current-based exponential synapses (CUBA) in BrainPy. For each presynaptic neuron *j* with spike train 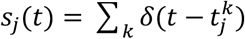, the synaptic state *x*_*j*_(*t*) followed:

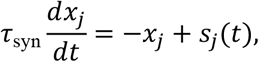

and contributed to the postsynaptic current as:

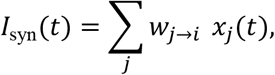

where *w*_*j*→*i*_ is the signed synaptic weight (positive for excitation and negative for inhibition). Unless otherwise stated, we used *τ*_exc_ = 5 ms with *w*_exc_ = 0.6, and *τ*_inh_ = 10 ms with *w*_inh_ = −6.7. Connectivity was instantiated as event-driven sparse projections with fixed connection probabilities obtained from the connectome-derived matrices. Conduction delays were set to 0.

The number of excitatory and inhibitory neurons in the network is scaled according to the anatomically derived neuron counts from each zebrafish brain region.^18^ To simulate ablation of a neuron class/morphotype in the SNN, we disconnected the targeted population by setting all incoming and outgoing connections involving that population to zero (i.e., *P*_*i*→*j*_ = 0 whenever *i* or *j* belongs to the ablated set). This operation removes the functional contribution of the targeted neurons without altering the dynamics of the remaining network:

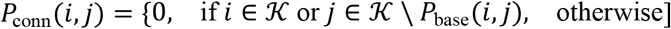

where *P*_conn_(*i, j*) is the connection probability between presynaptic neuron *i* and postsynaptic neuron *j, P*_base_(*i, j*) is the baseline (non-zero) probability determined by the connectome, and 𝒦 is the set of neurons targeted for ablation.

#### TIN reservoir construction

We constructed a zebrafish TIN reservoir network, where the connection weights were determined by the estimated number of potential synapses between neurons based on anatomical connectivity. Excitatory neurons were assigned positive weights, while inhibitory neurons were assigned negative weights. To ensure the reservoir dynamics remain in the echo state regime, we constrained the spectral radius *ρ*(*W*) of the recurrent weight matrix *W* to approximately 1:

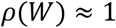

#### Conversion of calcium fluorescence signals into spike trains

To drive the spiking neural network with biologically realistic inputs, we converted experimentally recorded calcium fluorescence signals of retinal ganglion cells (RGCs) into input currents for tectal neurons (TNs) through a multi-step processing pipeline. First, calcium signals were deconvolved into estimated spike trains using a calcium dynamics-based inversion algorithm. The calcium dynamics were modeled as:

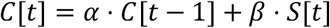

where *C*[*t*] represents the calcium concentration at time *t, S*[*t*] is the spike train, *α* = exp (−*dt*/*τ*) is the calcium decay constant with time constant *τ* = 0.5 s, and *β* = *dt*/*τ* is the calcium influx per spike. The spike train was estimated by inverting this relationship:

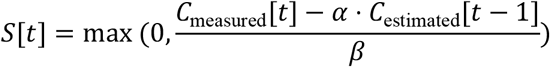

where *C*_measured_[*t*] is the experimentally recorded calcium signal and *C*_estimated_[*t*] is the estimated calcium concentration. The resulting spike trains were then converted to instantaneous firing rates using Gaussian temporal smoothing:

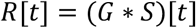

where *G* is a Gaussian kernel with standard deviation *σ* = 0.1 s (equivalent to a 100-ms smoothing window), and ∗ denotes convolution.

The firing rates were transformed into input currents using a linear gain relationship:

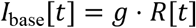

where *g* = 1 pA/(spikes/s) is the current gain factor.

Finally, the currents were interpolated to the simulation time base (0.1 ms resolution) using linear interpolation and scaled with channel-specific gain coefficients:

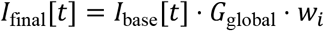

where *G*_global_ = 65 is the global gain coefficient, and *w*_*j*_ are channel-specific weights.

#### *In silico* cumulative and specific ablation of TIN morphotypes

We assessed morphotype contributions to visuomotor accuracy using two complementary procedures: (i) cumulative ablation to obtain an importance ranking, and (ii) specific ablation to further assess top-ranked morphotypes. Let ℳdenote the set of 44 TIN morphotypes. In cumulative ablation, we iteratively removed one morphotype at a time while keeping previously removed morphotypes disconnected. At iteration *k* with remaining set ℳ_*k*_, we computed accuracy for every candidate removal *m* ∈ ℳ_*k*_by running the network with ℳ_*k*_ \ {*m*} disconnected. We then removed the morphotype whose deletion caused the smallest accuracy reduction, thereby ranking morphotypes from least to most critical across iterations:

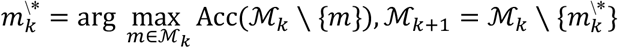

This procedure yields an ordered sequence 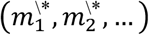 from least to most critical, where the last remaining/last removed morphotypes are the most important for maintaining accuracy. For specific ablation, we disconnected selected morphotypes (e.g., the top-ranked candidates) in the full network and quantified the resulting accuracy change relative to baseline.

#### Reservoir-specific dynamical analyses

To test whether the TIN network exhibits reservoir-like dynamics, we analyzed the same visual-input-driven TIN reservoir states used in the SNN simulations. For perturbation-decay analysis, each visual sequence was simulated twice with identical input: once from a zero initial state and once from a small random perturbation of the initial reservoir state. The Euclidean distance between the two state trajectories was normalized to the initial distance and averaged across visual sequences; progressive but stable decay indicated that perturbation-induced state differences were transiently preserved under input-driven dynamics. For fading-memory analysis, L2-regularized linear ridge readouts with standardized features were trained to reconstruct past RGC input vectors from current reservoir states across temporal delays using GroupKFold cross-validation across trial groups. Performance was quantified as cross-validated R^2. ER random and BA scale-free topology controls for these analyses were generated using the corresponding control recurrent networks, and were evaluated at the native recurrent weight scale rather than independently normalized to a common spectral radius. The no-recurrent control retained the same RGC-to-TIN input drive but replaced the recurrent TIN-to-TIN weight matrix with a zero matrix.

#### Signal-to-noise ratio quantification

To quantitatively assess the robustness of visuomotor transformation under noisy conditions, we calculated the signal-to-noise ratio (SNR) of TPN responses. The SNR was computed separately for the escape pathway (TPN-E population in response to looming stimuli) and the orienting pathway (TPN-O population in response to SMD stimuli). The response signal was defined as the total output of the relevant TPN population during stimulus presentation. The noise was defined as the baseline activity of the same TPN population over the matched pre-stimulus window. Specifically, for a given stimulus and its corresponding TPN population, the SNR was calculated as:

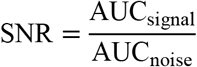

Where: AUC_signal_ is the area under the curve (AUC) of the summed firing rate of the relevant TPN population (either TPN-E or TPN-O) during the 15-second stimulus presentation period. AUC_noise_ is the AUC of the summed firing rate of the same TPN population during a 15-second pre-stimulus baseline period immediately preceding the stimulus onset. This baseline period corresponds to the initial 15 seconds of dynamic noise presentation in noisy trials.

#### 5-HT neuromodulation in the SNN model

To model serotonergic modulation, we implemented a connectivity-dependent increase in neuronal excitability for tectal neurons anatomically contacted with OT-projecting 5-HT axons. For each neuron *i*, we obtained a putative serotonergic connectivity score *p*_*i*_ (connection probability inferred by the same proximity-based procedure as above). During activation of a given 5-HT neuron subtype (DL_5-HT or SL_5-HT), we modified the spike threshold of neuron *i* as

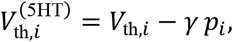

where *γ* controls the maximal modulation strength. This implementation increases the probability of spiking for neurons with stronger putative serotonergic innervation, capturing the experimentally observed increase in spontaneous firing following bath 5-HT application.

To quantify output bias under ambiguous sensory conditions (e.g., BMD), we computed

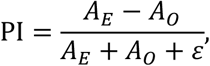

where *A*_*E*_ and *A*_*O*_ are the AUC responses of TPN-E and TPN-O populations over the standardized analysis window. Positive PI indicates escape bias, whereas negative PI indicates orienting bias.

#### Visual stimulation

All visual stimuli were presented monocularly to the left or right eye of the larva using a mini-projector (1920 × 1440 resolution) positioned laterally. A red filter was placed in front of the projector to minimize interference with fluorescence imaging. The stimulus protocols were as follows.

Looming stimulus: A black disk expanded symmetrically from 2° to whole screen of the visual field over 5 seconds on a bright background at a constant angular velocity, mimicking a high-risk threat. This was followed by a 10-second period where the screen gradually returned from black to the bright background. Each larva was tested with 5 trials, with an inter-trial interval of 120 seconds.

Moving dot stimuli: A single black dot moved horizontally across a white background in a reciprocating motion. The dot subtended either 4° (small moving dot, SMD, simulating prey), 8°,10°, or 16° (big moving dot, BMD, neutral stimulus) in diameter, depending on the experiment. Specifically, 8° BMD was used for *in vivo* calcium imaging of DL_5-HT and SL_5-HT neurons in Figure 5C, whereas 10° and 16° BMDs were used in the model-bias analyses and supplementary simulations as indicated in the corresponding figures. The dot completed three uninterrupted back-and-forth cycles over a 15-second period. Thus, the 15-second analysis window contained three continuous reciprocating cycles for moving-dot stimuli. Each larva received 5 trials per stimulus type, with a 120-second inter-trial interval. We used a 15-s analysis window across stimulus classes to standardize AUC-based quantification and model-input construction while preserving the full temporal extent of each stimulus epoch.

Noise stimuli: For experiments under degraded visual conditions, dynamic noise was superimposed on visual stimuli. The noise consisted of a field of white squares (3 × 3 pixels each), randomly repositioned at the frame rate of the projector. The noise level was defined as the total area of all noise squares divided by the total screen area. The noise presentation was temporally aligned with the stimulus: it began 15 seconds before the onset of the looming or moving dot stimuli, persisted throughout the 15-second stimulus duration, and continued for 15 seconds after the stimulus ended, resulting in a total noise duration of 45 seconds.

Light-dark transition: To assess the visual responses of 5-HT neurons, the background illumination was switched between bright (1.23 mW) and dark (0.167 mW) states. Each illumination period lasted 1 minute, and this cycle was repeated three times for a total of 6 minutes.

#### *In vivo* electrophysiological recording

Larvae were paralyzed with 100 μg/ml α-bungarotoxin (Sigma) for 10 - 15 min, then embedded in 2% low-melting point agarose. In whole-cell recording experiments, micropipettes were filled with an internal solution, which consisted of (in mM): 110 K-gluconate, 6 NaCl, 2 MgCl2, 2 CaCl2, 10 HEPES, and 10 EGTA (pH 7.41, 285 mOsm). The calculated chloride reversal potential (E_Cl_-) was ~ −60 mV. For TN whole-cell recording paired with calcium imaging in *Tg(elavl3:Hsa*.*H2B-GCaMP8f)*, EGTA was replaced with phosphocreatine to avoid calcium chelation. Cell-attached recording of TNs with bath application of 5-HT solution (dissolved in DMSO, 10 mM for stock solution) was performed in wild-type larvae.

For *in vivo* whole-cell recording, a small incision was made in the skin between the tectum and hindbrain using a glass pipette (~1 μm tip) after paralysis. After filling with the internal solution, the recording micropipette (20 - 25 MΩ in resistance, 1-mm tip opening) was advanced into the brain through the dissected hole and slowly approached the target cell with a persistent positive pressure to keep tip clean. After the contact of micropipette tip with TNs membrane, giga-ohm seal was formed by removing the positive pressure and applying a slight negative pressure. Whole-cell recording was achieved by delivering a few brief electrical zaps (50 μs to 2 ms) to break the cell membrane beneath the micropipette tip. Recordings were made with a patch-clamp amplifier (MultiClamp 700B, Axon Instruments), and signals were filtered at 5 kHz and sampled at 10 kHz by using Clampex 10.2 (Molecular Devices).

#### *In vivo* calcium imaging

Imaging was performed at room temperature with an Olympus Fluoview FVMPE-RS Multiphoton Laser Scanning Microscope equipped with a 20X water-immersion objective (1.0 NA). A Galvano scanning mode was used to achieve a frame rate of 4 - 5 Hz. For imaging RGC axonal arbors, *Tg(-7atoh7:GAL4-VP16);Tg(UAS:GCaMP6s)* larvae were used. For imaging the axonal arborizations of OT-projecting 5-HT neurons, *Ki(tph2:GAL4FF,myl7:EGFP);Tg(UAS:GCaMP6s)* larvae were used. GCaMP6s was excited at 920 nm. Image stacks were acquired with a voxel size of approximately 0.49 × 0.49 × 2.5 μm (x, y, z).

#### Calcium activity analysis

Acquired image stacks were first aligned to a reference volume using the MultiStackReg plugin in ImageJ to correct for motion artifacts. Regions of interest (ROIs) were manually defined based on distinct morphological features and regional signal activity. Calcium signals were extracted as the mean fluorescence intensity within each ROI over time. The fluorescence change (ΔF/F_0_) was computed using a custom Python script. The baseline fluorescence (F_0_) was defined as the average fluorescence intensity during a 10-second window preceding stimulus onset. The ΔF/F_0_ was calculated as (F - F_0_)/F_0_.

#### Ablation of OT-projecting 5-HT neurons

Targeted ablations were performed on an Olympus Fluoview FVMPE-RS multiphoton microscope using a 20×/1.0 NA water-immersion objective. 6-8 dpf larvae of *Ki(tph2:GAL4FF,myl7:EGFP);Tg(5xUAS-hsp70l:mScarlet);Tg2(elavl3:GCaMP6s)* were used to identify OT-projecting pretectal 5-HT neurons. Pretectal 5-HT neurons were classified by their distinct visual response patterns: SL_5-HT neurons were defined by sustained ON responses to bright background illumination, whereas DL_5-HT neurons were defined by transient ON and OFF responses to illumination changes.

Single-cell ablation was achieved by focused 800-nm laser scanning targeted to the soma and primary neurite until fluorescence was irreversibly extinguished. Successful ablation was verified by loss of mScarlet signal and absence of stimulus-evoked calcium activity in the targeted cell. Larvae were then released from agarose and allowed to recover in facility water for ≥18 h before post-ablation imaging and behavioral assays. Animals with incomplete ablation or unintended damage were excluded from analysis.

#### Assay for BMD-evoked behaviors

Larval zebrafish (6 - 8 dpf) were placed individually in a custom behavioral arena with a vertical projection screen displaying visual stimuli. Behavioral responses to BMD stimuli (10° in diameter) were recorded using a high-speed, infrared-sensitive camera (Redlake Motionscope M3) at 30 frames per second. Each larva was tested in 10 trials with a 120-second inter-trial interval. We used the fish-to-screen distance during BMD presentation as a behavioral proxy for orienting *versus* escape bias. A decrease in distance indicates approach toward the stimulus-bearing screen and was interpreted as orienting, whereas an increase in distance indicates movement away from the screen and was interpreted as escape bias. This metric is useful because BMD is an ambiguous stimulus that does not reliably trigger a single stereotyped motor program.

For automated tracking and distance quantification, zebrafish position was automatically tracked throughout each trial using a custom Python analysis pipeline. The analysis involved three main computational steps. (1) Screen reference calibration: for each experimental session, a screen reference line was manually defined by selecting two points along the projection screen boundary in the first frame. This line served as the spatial reference for all distance calculations. (2) Fish position detection: fish position was detected in each frame using an advanced background subtraction algorithm. A background model was first constructed by median-averaging 50 randomly sampled frames. For each subsequent frame, the absolute difference from the background model was computed, thresholded, and processed with morphological operations to reduce noise. The largest valid contour (50 - 2000 pixel area) was identified, and its centroid was calculated as the fish’s position. (3) Distance to screen calculation: the Euclidean distance between the detected fish position and the screen reference line was computed using the point-to-line distance formula:

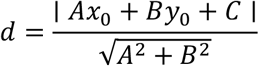

where (*x*_0_, *y*_0_) represents the fish coordinates, and *A, B, C* are the coefficients of the line equation defined by the two reference points. Trajectories were smoothed using a 5-frame moving average filter to reduce high-frequency noise.

### QUANTIFICATION AND STATISTICAL ANALYSIS

Data presentation and software: Data are presented as mean ± standard error of the mean (SEM) throughout the manuscript, unless otherwise specified in the figure legends. Unless otherwise stated, line traces and bar plots show the mean, and error bars indicate SEM. Repeated trials from the same biological sample (for example, multiple trials from the same larva, or repeated recordings from the same neuron before and during bath application) were averaged within sample before statistical testing. For behavioral assays with repeated trials from the same larva, statistical tests were performed on per-larva means even when individual trials are displayed in the figures. In *in silico* modeling, n refers to the number of independent simulation trials; in *in vivo* experiments, n refers to the number of biologically independent samples, unless otherwise specified in the figure legends. Two-sided Wilcoxon signed-rank tests were used for paired comparisons. Two-sided Mann–Whitney U tests were used for comparisons between two independent groups. For quantities tested against a theoretical null value (for example, a fold-change relative to 1), two-sided one-sample Wilcoxon signed-rank tests were used. Exact n values, statistical tests, and definitions of graphical summaries are provided in the corresponding figure legends. All statistical analyses were performed using custom scripts in Python.

Normality testing and test selection: The choice of parametric or non-parametric statistical tests was based on the nature and distribution of the data. For comparisons of data derived from *in silico* simulations (e.g., accuracy and robustness across trials) and other datasets where the assumption of normality could not be guaranteed, we used non-parametric tests. The two-sided Wilcoxon signed-rank test was employed for paired comparisons (e.g., comparing network performance with *vs*. without TINs across multiple simulation trials), as it does not assume a normal distribution and is robust to outliers.

Randomization and blinding: For *in vivo* experiments involving group comparisons (e.g., 5-HT neuron ablation), larvae were randomly assigned to experimental groups. Due to the necessity of targeting fluorescently labeled neurons or structures, blinding was not always feasible during data acquisition. However, automated analysis pipelines (for behavior and calcium imaging) and objective criteria (for electrophysiology and model output) were used for quantification to minimize potential bias.

Significance and precision: Statistical significance was defined as *P* < 0.05. Exact *P*-values are reported in the figures or figure legends, or denoted as follows: **\****P* < 0.05, **\*\****P* < 0.01, **\*\*\****P* < 0.001.

