## Supplementary Figures and Legends for "A tectal reservoir implements adaptive visuomotor transformation via serotonergically coordinated push-pull-like mechanisms"

### SUPPLEMENTAL FIGURE AND LEGENDS

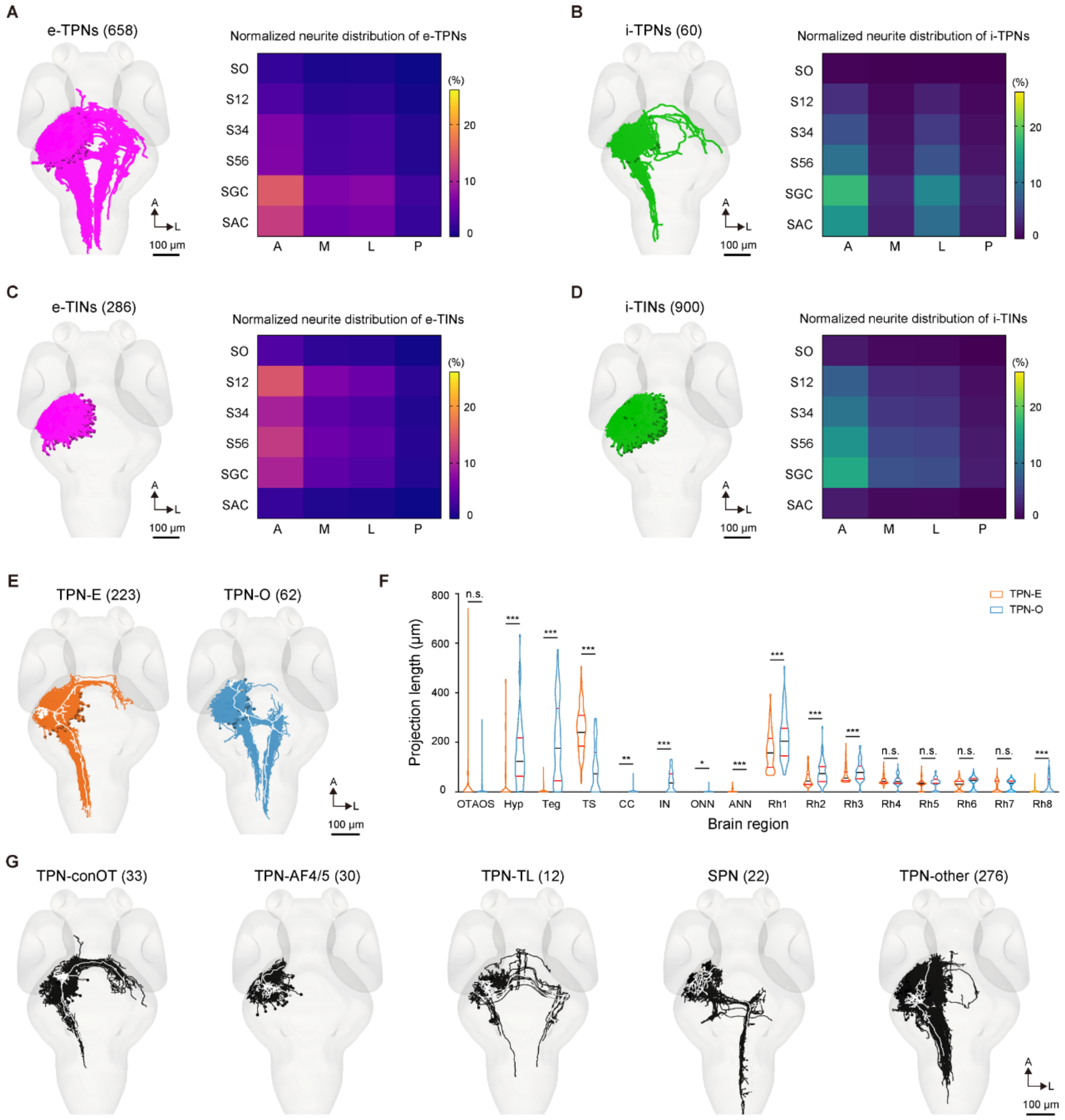


**Figure S1. Classification of tectal neurons and excitatory tectal projection neurons, related to Figure 1.**

(A, B) Morphology (left) and normalized neurite distribution within the OT neuropil (right) of tectal projection neurons (TPNs), including excitatory TPNs (e-TPNs, A) and inhibitory TPNs (i-TPNs, B).

(C, D) Morphology (left) and normalized neurite distribution within the OT neuropil (right) of tectal interneurons (TINs), including excitatory TINs (e-TINs, C) and inhibitory TINs (i-TINs, D).

(E, F) Morphology (E) and distinct innervation patterns (F) of TPN-E and TPN-O neurons. OTAOS, optic tract and accessory optic system; Hyp, hypothalamus; Teg, tegmentum; TS, torus semicircularis; CC, corpus cerebelli; IN, interpeduncular nucleus; ONN, oculomotor nerve nucleus; ANN, abducens nerve nucleus; Rh1-8, rhombomere 1-8. The internal black line represents the median, and the red lines indicate the 25th and 75th percentiles. n.s., no significance; *$P < 0.05$; **$P < 0.01$; ***$P < 0.001$ (Two-sided Mann-Whitney U tests).White traces show representative neuron morphologies.

(G) Other subtypes of e-TPNs classified by downstream projection targets or soma location. TPN-ConOT, e-TPNs projecting to contralateral OT; TPN-AF4/5, e-TPNs projecting to the arborization fields 4 and 5 (AF4 and AF5); TPN-TL, e-TPNs projecting to the torus longitudinalis (TL); SPN, e-TPNs with soma at the neuropil; TPN-other, e-TPNs projecting to other targets. White traces show representative neuron morphologies.

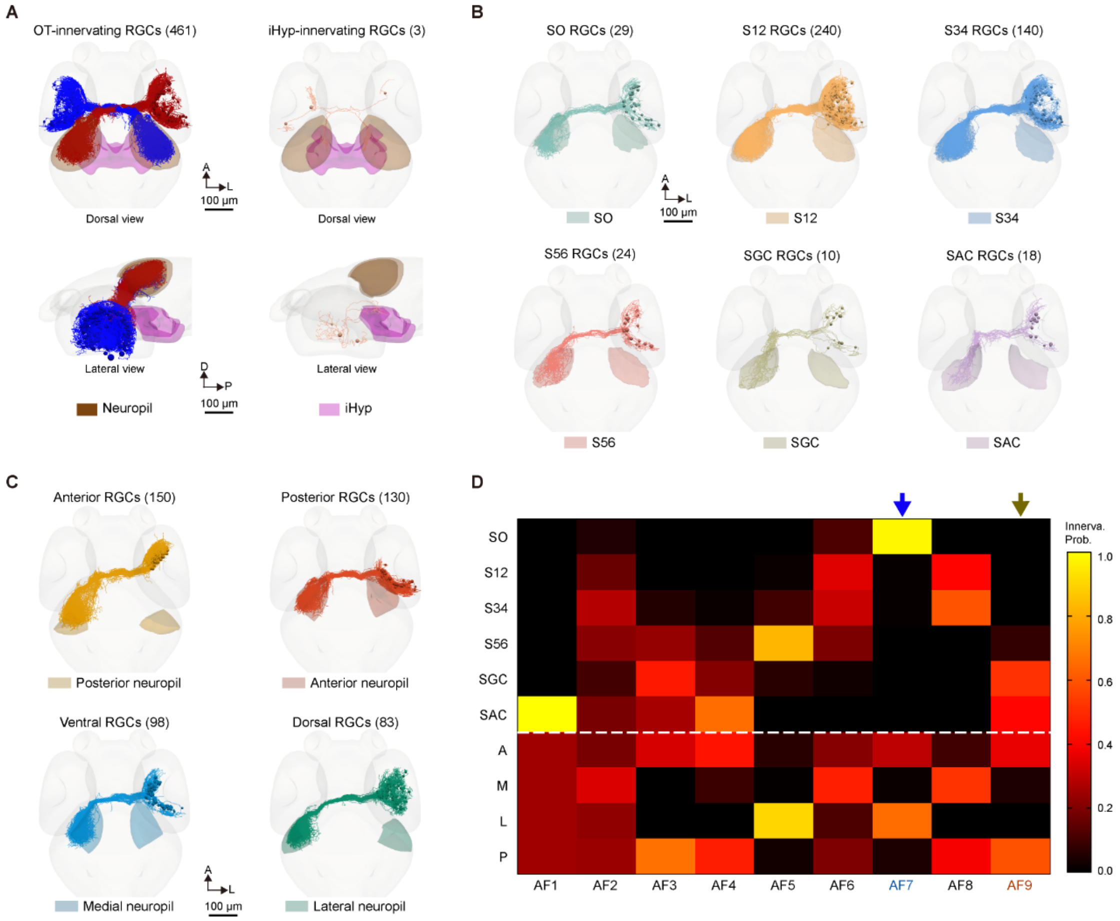


**Figure S2. Laminar and topographic projections of RGCs, related to Figure 1.**

(A) Classification of RGCs by projection targets. Most RGCs (461/464, left) project to the contralateral OT, and a minority of RGCs (3/464, right) project to the intermediate hypothalamus (iHyp). Top, dorsal view; bottom, lateral view.

(B, C) Laminar (B) and topographic (C) specificity of RGC axonal arborizations in the OT neuropil.

(D) Quantitative mapping of AF-projecting RGC axonal innervation in the OT neuropil using normalized axonal lengths across 6 neuropil layers and 4 topographic zones. For each AF-projecting RGC, the cumulative projection weights in the six neuropil layers (above the dashed line) and the four spatial columns (below the dashed line) are summed to unity. The blue and brown arrows represent the axonal innervation patterns of AF7- and AF9-projecting RGCs, respectively.

Numbers in parentheses indicate the number of cells analyzed. AF, arborization field; A, anterior; D, dorsal; L, lateral; P, posterior.

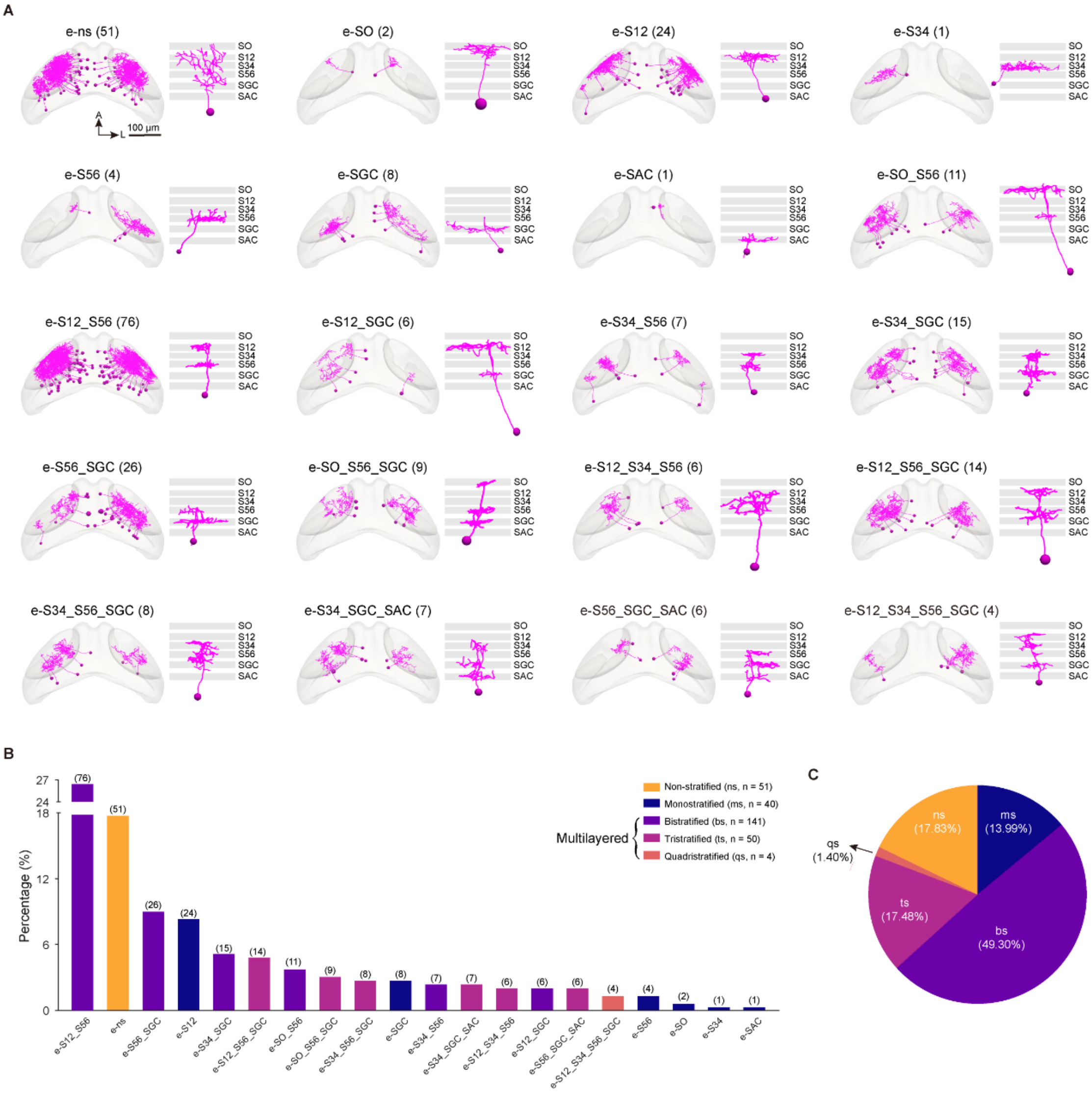
A
e-ns (51)
e-SO (2)
e-S12 (24)
e-S34 (1)
e-S56 (4)
e-SGC (8)
e-SAC (1)
e-SO_S56 (11)
e-S12_S56 (76)
e-S12_SGC (6)
e-S34_S56 (7)
e-S34_SGC (15)
e-S56_SGC (26)
e-SO_S56_SGC (9)
e-S12_S34_S56 (6)
e-S12_S56_SGC (14)
e-S34_S56_SGC (8)
e-S34_SGC_SAC (7)
e-S56_SGC_SAC (6)
e-S12_S34_S56_SGC (4)
SO
S12
S34
S56
SGC
SAC
A
L
100 µm
B
Percentage (%)
27
24
18
12
6
0
(76)
(51)
(26)
(24)
(15)
(14)
(11)
(9)
(8)
(8)
(7)
(7)
(6)
(6)
(6)
(4)
(4)
(2)
(1)
(1)
e-S12_S56
e-ns
e-S56_SGC
e-S12
e-S34_SGC
e-S12_S56_SGC
e-SO_S56
e-SO_S56_SGC
e-S34_S56_SGC
e-SGC
e-S34_S56
e-S34_SGC_SAC
e-S12_S34_S56
e-S12_SGC
e-S56_SGC_SAC
e-S12_S34_S56_SGC
e-S56
e-SO
e-S34
e-SAC
Non-stratified (ns, n = 51)
Monostratified (ms, n = 40)
Multilayered
Bistratified (bs, n = 141)
Tristratified (ts, n = 50)
Quadristratified (qs, n = 4)
C
ns
(17.83%)
ms
(13.99%)
qs
(1.40%)
ts
(17.48%)
bs
(49.30%)

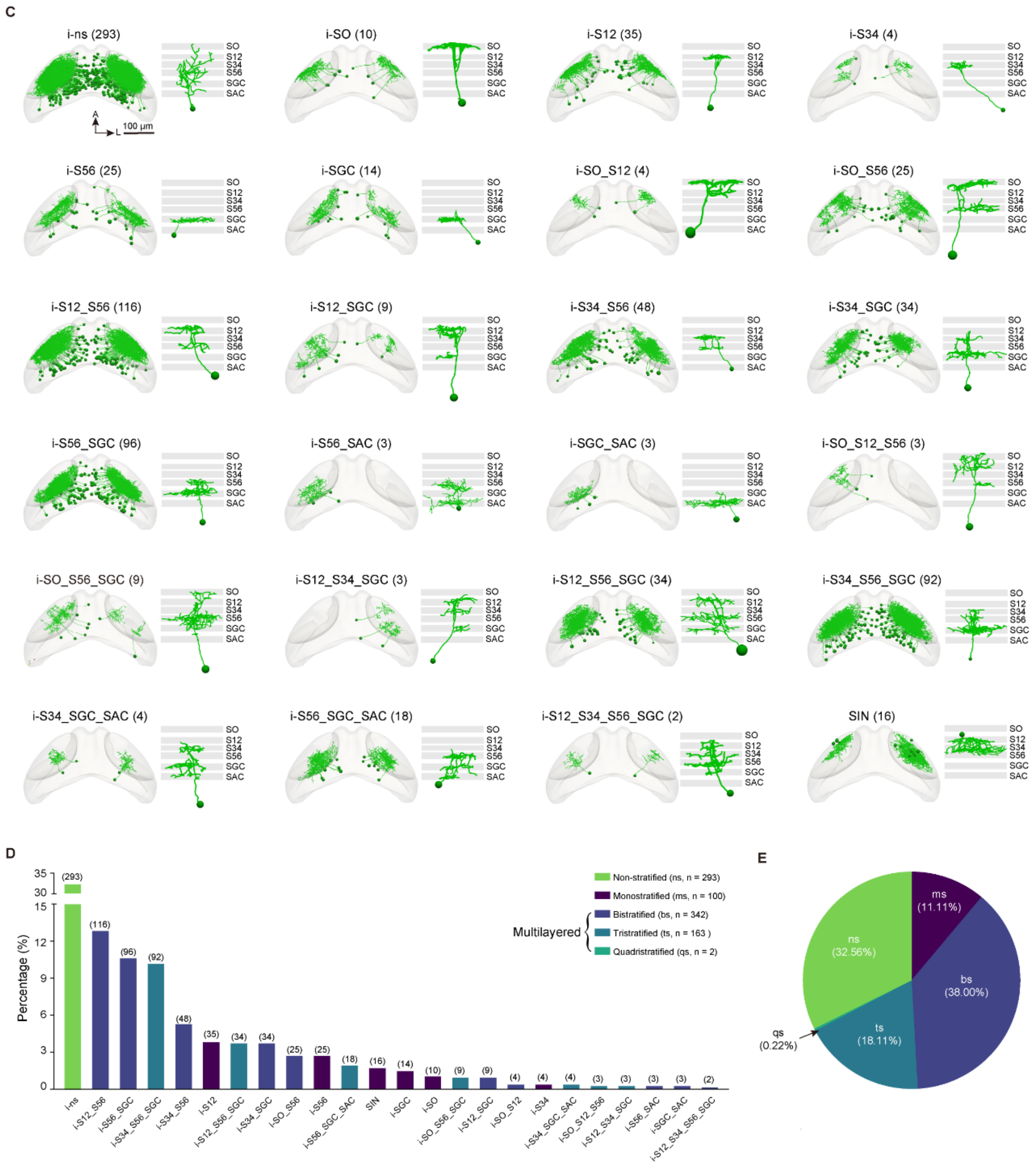


**Figure S3. Excitatory and inhibitory morphotypes of TINs, related to Figure 2.**

(A) Twenty e-TIN morphotypes classified by their neurite distribution across six laminae in the OT neuropil.

(B) Percentages of 20 e-TIN morphotypes, which can be classified into 5 categories (color coded): non-stratified (ns), monostratified (ms), bistratified (bs), tristratified (ts), and quadristratified (qs) neurites.

(C) Pie chart summarizing the distribution of 5 e-TIN categories, color-coded according to neurite stratification patterns.

(D) Twenty-four morphotypes of i-TINs classified by their neurite distribution across the six laminae of the OT neuropil.

(E) Percentages of 24 i-TIN morphotypes, classified into 5 categories based on their neurite stratification: non-stratified (ns), monostratified (ms), bistratified (bs), tristratified (ts), and quadristratified (qs) neurites.

(F) Pie plot of 5 i-TIN categories.

A, anterior; L, lateral. Numbers in parentheses (A, B, D, E) indicate the cell number. Numbers in the parentheses (C, F) indicate the percentages.

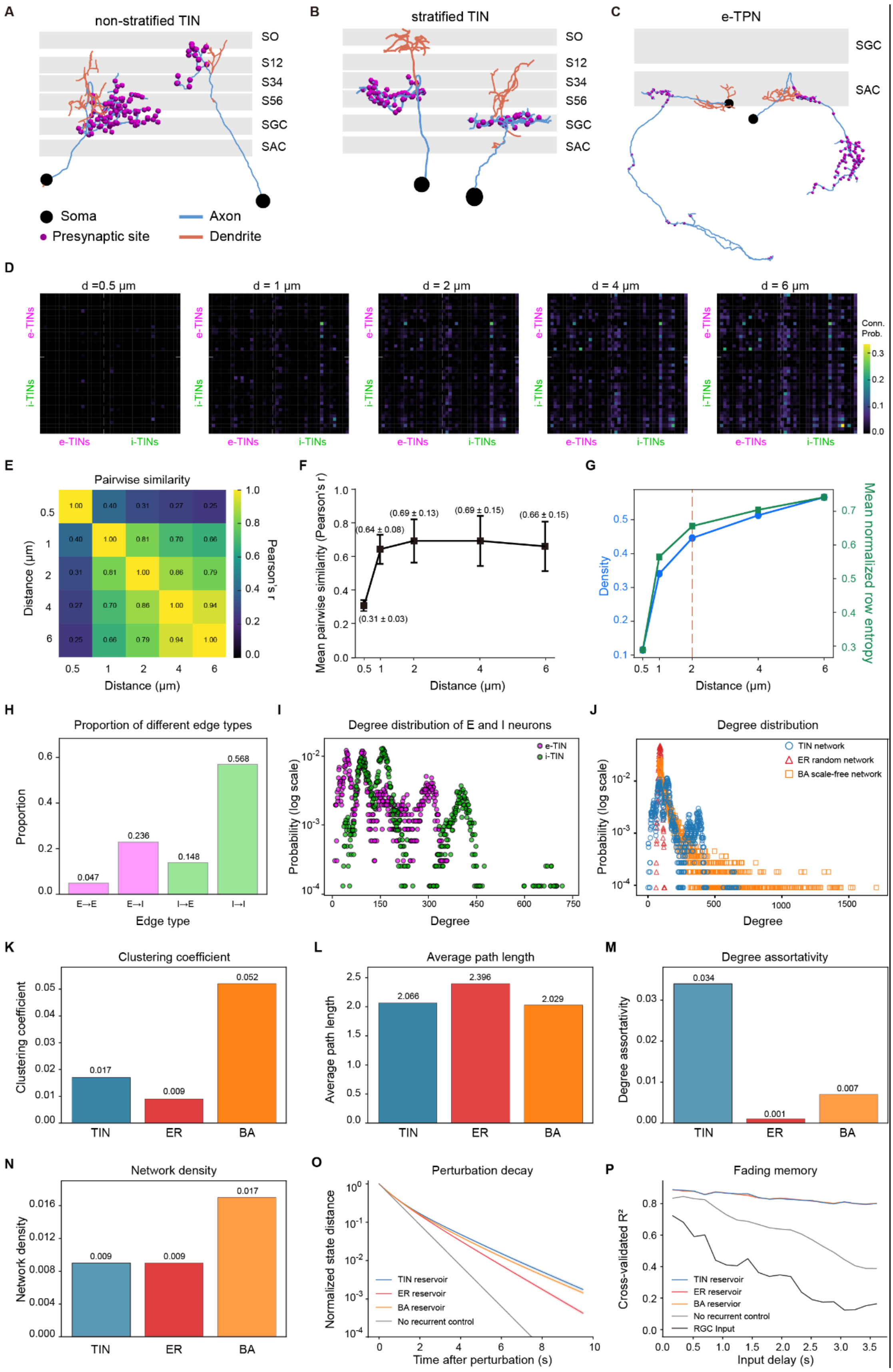

A
non-stratified TIN
SO
S12
S34
S56
SGC
SAC
Soma
Presynaptic site
Axon
Dendrite
B
stratified TIN
C
e-TPN
D
d =0.5 μm
d = 1 μm
d = 2 μm
d = 4 μm
d = 6 μm
e-TINs
i-TINs
Conn. Prob.
E
Pairwise similarity
Distance (μm)
Pearson's r
F
Mean pairwise similarity (Pearson's r)
(0.31 ± 0.03)
(0.64 ± 0.08)
(0.69 ± 0.13)
(0.69 ± 0.15)
(0.66 ± 0.15)
G
Density
Mean normalized row entropy
H
Proportion of different edge types
Proportion
Edge type
E→E
E→I
I→E
I→I
I
Degree distribution of E and I neurons
e-TIN
i-TIN
Probability (log scale)
Degree
J
Degree distribution
TIN network
ER random network
BA scale-free network
K
Clustering coefficient
TIN
ER
BA
L
Average path length
M
Degree assortativity
N
Network density
O
Perturbation decay
Normalized state distance
TIN reservoir
ER reservoir
BA reservoir
No recurrent control
Time after perturbation (s)
P
Fading memory
Cross-validated R²
RGC Input
Input delay (s)

**Figure S4. Axon-dendrite discrimination of TNs, and the connectivity thresholding, network architecture, and performance of the TIN reservoir, related to Figure 2.**

(A) Morphology of two non-stratified TINs. Dendrites and axons intermingle across multiple OT neuropil layers without clear laminar restriction.

(B) Morphology of two stratified TINs. Dendrites are confined to relatively superficial neuropil layers, whereas axons arborize in relatively deeper layers.

(C) Morphology of two excitatory tectal projection neurons (e-TPNs). Dendrites are localized within the OT neuropil, whereas the axon exits the OT. The presynaptic puncta (purple dots) on these neurons' axonal arbors were labeled with the presynaptic marker Synaptobrevin-GFP (Sypb-EGFP) (see STAR Methods). Dendritic (orange), axonal (blue), and presynaptic (purple) sites are shown for each neuron. Layered regions of the OT neuropil are indicated. The soma of each neuron is marked by a black dot.

(D) Morphotype-level connectivity matrices generated using axon-dendrite distances of 0.5, 1, 2, 4, and 6 μm. Each matrix entry indicates the directed connection probability between source (x-axis) and target (y-axis) morphotypes, inferred from connectome-derived putative anatomical connectivity (see STAR methods).

(E) Pairwise similarity (Pearson's r) between morphotype-level connectivity matrices generated at different distances. Similarity was calculated between flattened directed connection probability matrices, with warmer colors indicating higher similarity.

(F) Mean pairwise similarity of morphotype-level connectivity matrices across distances. Values indicate mean ± SEM across pairwise Pearson's r comparisons between each matrix and the remaining matrices.

(G) Network density (blue; fraction of non-zero morphotype-to-morphotype entries) and mean row selectivity (green, see STAR Methods) across distances. Density increased monotonically, whereas row selectivity decreased with larger distances, consistent with progressively more diffuse connectivity estimates. The dashed line marks the 2 μm used throughout this study.

(H, I) Network characteristics of the TIN reservoir. Proportions of edge types (H), showing excitatory-to-excitatory (E→E), excitatory-to-inhibitory (E→I), inhibitory-to-excitatory (I→E), and inhibitory-to-inhibitory (I→I) connections. Degree distribution of excitatory and inhibitory TINs (I), showing the number of incoming and outgoing connections per neuron.

(J-N) Comparison of network properties between the TIN reservoir (blue), an Erdős–Rényi (ER) random network (red), and a Barabási-Albert (BA) scale-free network (orange): degree distribution (J), clustering coefficient (K), average path length (L), degree assortativity (M), and network density (N).

(O, P) Reservoir-state dynamics of the TIN network. (O) Perturbation-decay analysis comparing the TIN reservoir (blue), ER random reservoir (red), BA scale-free reservoir (orange), and no-recurrent control (gray). Each visual sequence was simulated from a zero initial state and from a small random perturbation under identical RGC drives; curves show the normalized Euclidean distance between the two reservoir-state trajectories over time. (P) Fading-memory analysis. L2-regularized linear ridge readouts reconstructed recent RGC input history from current reservoir states across temporal delays, with performance quantified as cross-validated R2. The RGC input trace (black) provides an input baseline. ER and BA topology controls were evaluated at their native recurrent weight scale rather than independently normalized to a common spectral radius, and the no-recurrent control retained the same RGC-to-TIN input drive while setting all recurrent TIN-to-TIN weights to zero.

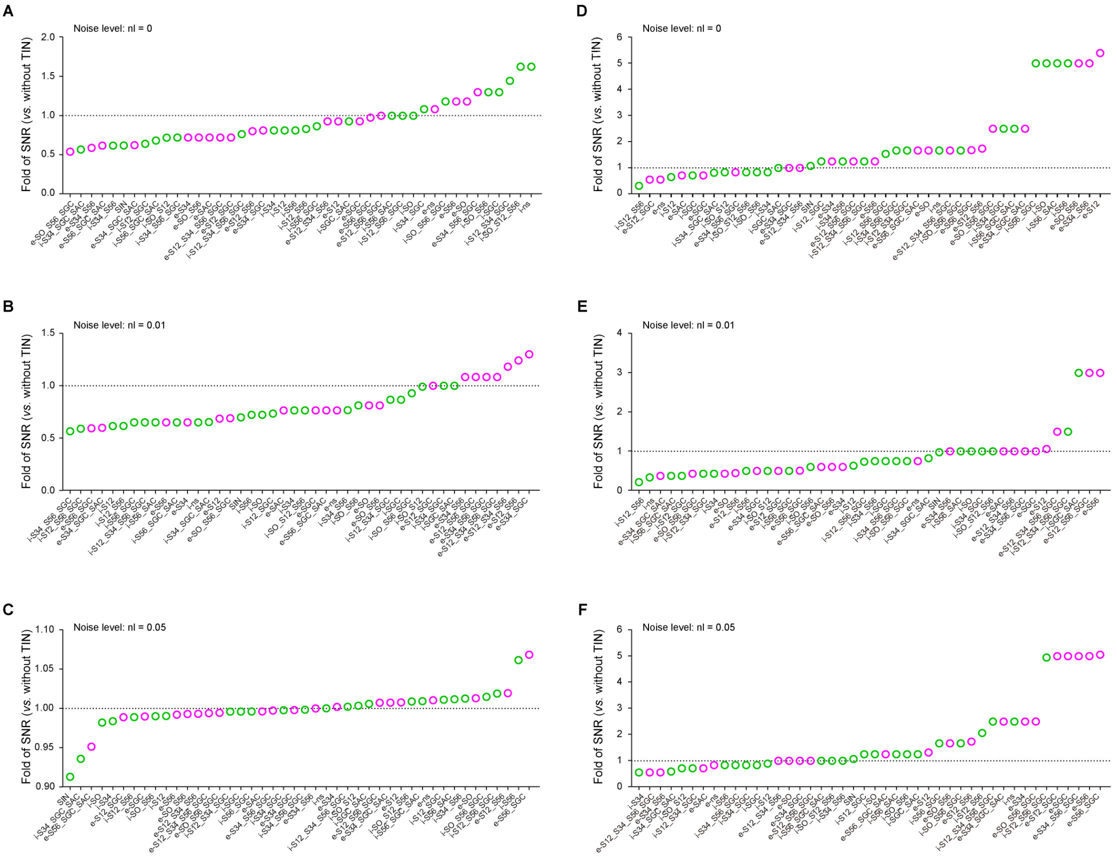


**Figure S5. Morphotype-specific contributions of TINs to looming- and SMD-evoked response robustness across no- and low-noise conditions, related to Figure 3.**

(A-C) Fold-change in looming-evoked TPN-E SNR relative to the model without the TIN network for each TIN morphotype activation at noise levels nl = 0 (A), 0.01 (B), and 0.05 (C).

(D-F) Fold-change in SMD-evoked TPN-O SNR relative to the model without the TIN network for each TIN morphotype activation at noise levels nl = 0 (D), 0.01 (E), and 0.05 (F).

Magenta circles denote e-TIN morphotypes and green circles denote i-TIN morphotypes. Morphotypes were ranked by their contribution at each noise level.

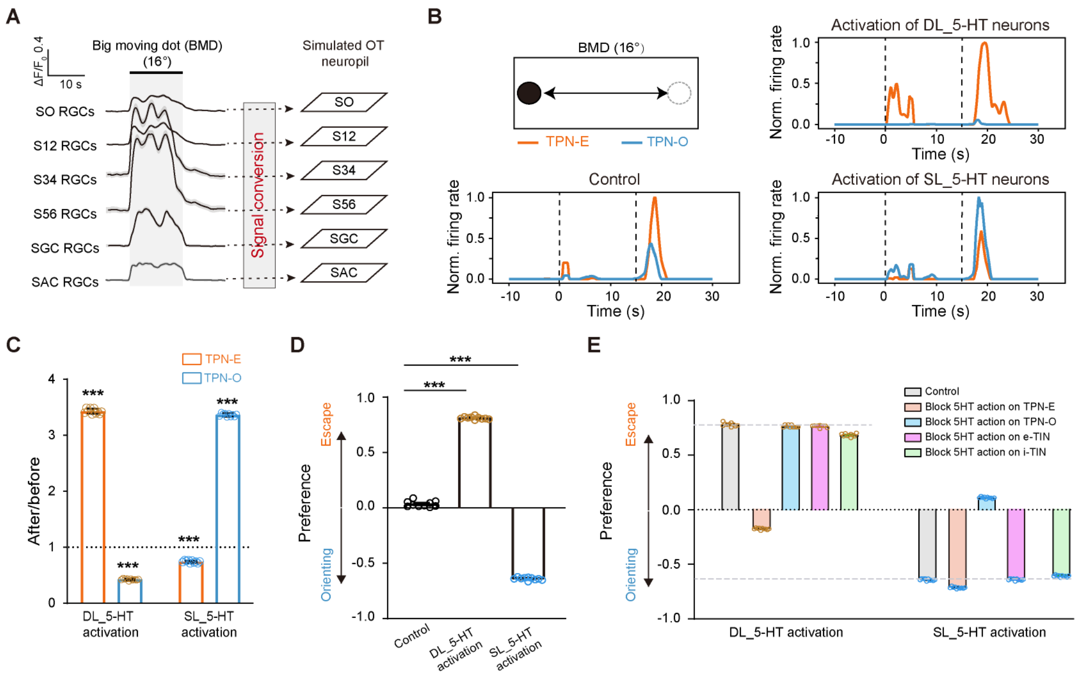


**Figure S6. DL_5-HT and SL_5-HT neurons bias model outputs under 16° BMD, related to Figure 4.**

(A) Layer-specific calcium activities of RGC axonal arbors in the OT neuropil, evoked by big moving dot (BMD, 16° in diameter). These calcium activities served as inputs to the OT SNN model after conversion into spike-proxy current inputs. Left, BMD-evoked RGC responses. Right, schematic showing that RGC activity is fed into the model. Activity traces represent the mean ± SEM averaged from 7 larvae (2 trials per larva); error bars are too small to be visible. The imaging was performed on double-transgenic *Tg(-7atoh7:GAL4-VP16);Tg(UAS:GCaMP6s)* larvae at 6-8 dpf.

(B) BMD-evoked TPN-E and TPN-O responses under control conditions without 5-HT neuronal activation (bottom left), with activation of DL_5-HT neurons (top right), or with activation of SL_5-HT neurons (bottom right). Top left, schematic of BMD stimulus, which lasts 15 s. The window between the two dashed lines indicates the visual stimulation period. Traces are averaged from 5 simulation trials.

(C) DL_5-HT and SL_5-HT neuronal activation-induced changes in 16° BMD-evoked TPN-E and TPN-O responses. Each point denotes one matched simulation trial (n = 10). The dashed line indicates the no-change level (fold-change = 1). ***$P$ < 0.001 (two-sided one-sample Wilcoxon signed-rank tests against 1 for each group).

(D) DL_5-HT and SL_5-HT neuronal activation-induced changes in 16° BMD-evoked behavioral preference. The preference index was calculated as (AUC of TPN-E minus AUC of TPN-O) divided by the sum of both AUCs. ***$P$ < 0.001 (two-sided Wilcoxon signed-rank tests comparing each 5-HT activation condition with control conditions).

(E) Component-wise blocking analysis of behavioral preference shifts induced by DL_5-HT or SL_5-HT neuronal activation, revealing differential contributions of OT neuron classes.
