## Supplementary Tables for "A tectal reservoir implements adaptive visuomotor transformation via serotonergically coordinated push-pull-like mechanisms"

[5] Lead contact

### SUPPLEMENTAL TABLES

**Table S1. Parameters for LIF neuron models of key cell types, related to Figure 1**

| | RGCs | TNs | 5-HT neurons |
|---|---|---|---|
| $\tau_m$ (membrane time constant, ms) | 5 - 20 | 5 - 20 | 15 - 30 |
| $V_{rest}$ (resting potential, mV) | -60 | -60 | -60 |
| $V_{th}$ (threshold, mV) | -50 to -55 | -50 to -55 | -50 to -55 |
| $V_{reset}$ (resetting potential, mV) | -60 to -65 | -60 to -65 | -60 to -65 |
| $R_m$ (membrane resistance, MΩ) | 100 - 200 | 100 - 200 | 100 - 150 |
| $\tau_{ref}$ (refractory period, ms) | 2 - 5 | 2 - 5 | 2 - 5 |

**Table S2. The connection probability between neuron types, related to Figures 1-3 (Part 1 of 4; from SO RGC to e-S12_S56 TIN).**

| | SO RGC | S12 RGC | S34 RGC | S56 RGC | SGC RGC | SAC RGC | e-SO TIN | e-SO_S56 TIN | e-SO_S56_SGC TIN | e-S12 TIN | e-S12_S34_S56 TIN | e-S12_S34_S56_SGC TIN | e-S12_S56 TIN |
|---|---|---|---|---|---|---|---|---|---|---|---|---|---|
| **SO RGC** | 0 | 0 | 0 | 0 | 0 | 0 | 0 | 0 | 0 | 0.025747 | 0 | 0 | 0 |
| **S12 RGC** | 0 | 0 | 0 | 0 | 0 | 0 | 0 | 0 | 0 | 0.033986 | 0 | 0 | 0.02111 |
| **S34 RGC** | 0 | 0 | 0 | 0 | 0 | 0 | 0 | 0 | 0 | 0.01131 | 0 | 0 | 0.030282 |
| **S56 RGC** | 0 | 0 | 0 | 0 | 0 | 0 | 0 | 0 | 0.00463 | 0 | 0 | 0 | 0 |
| **SGC RGC** | 0 | 0 | 0 | 0 | 0 | 0 | 0 | 0 | 0.011111 | 0 | 0 | 0 | 0.001316 |
| **SAC RGC** | 0 | 0 | 0 | 0 | 0 | 0 | 0 | 0 | 0 | 0 | 0 | 0 | 0 |
| **e-SO TIN** | 0 | 0 | 0 | 0 | 0 | 0 | 0 | 0 | 0 | 0 | 0 | 0 | 0 |
| **e-SO_S56 TIN** | 0 | 0 | 0 | 0 | 0 | 0 | 0 | 0 | 0.010101 | 0 | 0 | 0 | 0 |
| **e-SO_S56_SGC TIN** | 0 | 0 | 0 | 0 | 0 | 0 | 0 | 0.020202 | 0.024691 | 0 | 0 | 0 | 0.00731 |
| **e-S12 TIN** | 0 | 0 | 0 | 0 | 0 | 0 | 0 | 0 | 0 | 0.010417 | 0 | 0 | 0.000548 |
| **e-S12_S34_S56 TIN** | 0 | 0 | 0 | 0 | 0 | 0 | 0 | 0.030303 | 0 | 0 | 0 | 0 | 0.006579 |
| **e-S12_S34_S56_SGC TIN** | 0 | 0 | 0 | 0 | 0 | 0 | 0 | 0 | 0.037037 | 0.013889 | 0 | 0 | 0.008772 |
| **e-S12_S56 TIN** | 0 | 0 | 0 | 0 | 0 | 0 | 0 | 0.011962 | 0.00731 | 0.011513 | 0.002193 | 0 | 0 |
| **e-S12_S56_SGC TIN** | 0 | 0 | 0 | 0 | 0 | 0 | 0 | 0.038961 | 0.015873 | 0.011905 | 0 | 0 | 0.008459 |
| **e-S12_SGC TIN** | 0 | 0 | 0 | 0 | 0 | 0 | 0 | 0.015152 | 0 | 0.020833 | 0 | 0 | 0.004386 |
| **e-S34 TIN** | 0 | 0 | 0 | 0 | 0 | 0 | 0 | 0 | 0 | 0.041667 | 0 | 0 | 0.013158 |
| **e-S34_S56 TIN** | 0 | 0 | 0 | 0 | 0 | 0 | 0 | 0.012987 | 0 | 0 | 0 | 0 | 0.007519 |

| | SO RGC | S12 RGC | S34 RGC | S56 RGC | SGC RGC | SAC RGC | e-SO TIN | e-SO_S56 TIN | e-SO_S56_SGC TIN | e-S12 TIN | e-S12_S34_S56 TIN | e-S12_S34_S56_SGC TIN | e-S12_S56 TIN |
|---|---|---|---|---|---|---|---|---|---|---|---|---|---|
| **e-S34_S56_SGC TIN** | 0 | 0 | 0 | 0 | 0 | 0 | 0 | 0.022727 | 0.041667 | 0 | 0 | 0 | 0.011513 |
| **e-S34_SGC TIN** | 0 | 0 | 0 | 0 | 0 | 0 | 0 | 0.018182 | 0.02963 | 0 | 0 | 0 | 0.007018 |
| **e-S34_SGC_SAC TIN** | 0 | 0 | 0 | 0 | 0 | 0 | 0 | 0.012987 | 0 | 0 | 0 | 0 | 0.018797 |
| **e-S56 TIN** | 0 | 0 | 0 | 0 | 0 | 0 | 0 | 0 | 0 | 0 | 0 | 0 | 0 |
| **e-S56_SGC TIN** | 0 | 0 | 0 | 0 | 0 | 0 | 0 | 0 | 0.025641 | 0 | 0 | 0 | 0 |
| **e-S56_SGC_SAC TIN** | 0 | 0 | 0 | 0 | 0 | 0 | 0 | 0 | 0 | 0 | 0 | 0.066667 | 0 |
| **e-SGC TIN** | 0 | 0 | 0 | 0 | 0 | 0 | 0 | 0 | 0.041667 | 0 | 0 | 0 | 0 |
| **e-SAC TIN** | 0 | 0 | 0 | 0 | 0 | 0 | 0 | 0 | 0 | 0 | 0 | 0 | 0 |
| **e-ns TIN** | 0 | 0 | 0 | 0 | 0 | 0 | 0 | 0.00713 | 0.019608 | 0.008987 | 0.003268 | 0 | 0.007224 |
| **i-SO TIN** | 0 | 0 | 0 | 0 | 0 | 0 | 0 | 0.009091 | 0 | 0.004167 | 0 | 0 | 0.002597 |
| **i-SO_S12 TIN** | 0 | 0 | 0 | 0 | 0 | 0 | 0 | 0 | 0 | 0 | 0 | 0 | 0 |
| **i-SO_S12_S56 TIN** | 0 | 0 | 0 | 0 | 0 | 0 | 0 | 0 | 0 | 0 | 0 | 0 | 0 |
| **i-SO_S56 TIN** | 0 | 0 | 0 | 0 | 0 | 0 | 0 | 0 | 0 | 0.008333 | 0 | 0 | 0.004675 |
| **i-SO_S56_SGC TIN** | 0 | 0 | 0 | 0 | 0 | 0 | 0 | 0 | 0.012346 | 0.00463 | 0 | 0 | 0.007215 |
| **i-S12 TIN** | 0 | 0 | 0 | 0 | 0 | 0 | 0 | 0 | 0 | 0.005952 | 0 | 0 | 0.004325 |
| **i-S12_S34_S56_SGC TIN** | 0 | 0 | 0 | 0 | 0 | 0 | 0 | 0 | 0 | 0 | 0 | 0 | 0 |
| **i-S12_S34_SGC TIN** | 0 | 0 | 0 | 0 | 0 | 0 | 0 | 0 | 0 | 0 | 0 | 0 | 0 |
| **i-S12_S56 TIN** | 0 | 0 | 0 | 0 | 0 | 0 | 0 | 0 | 0.000958 | 0.007902 | 0 | 0 | 0.003135 |
| **i-S12_S56_SGC TIN** | 0 | 0 | 0 | 0 | 0 | 0 | 0 | 0 | 0.013072 | 0.008578 | 0 | 0 | 0.009931 |

| | SO RGC | S12 RGC | S34 RGC | S56 RGC | SGC RGC | SAC RGC | e-SO TIN | e-SO_S56 TIN | e-SO_S56_SGC TIN | e-S12 TIN | e-S12_S34_S56 TIN | e-S12_S34_S56_SGC TIN | e-S12_S56 TIN |
|---|---|---|---|---|---|---|---|---|---|---|---|---|---|
| **i-S12_SGC TIN** | 0 | 0 | 0 | 0 | 0 | 0 | 0 | 0 | 0.012346 | 0 | 0 | 0 | 0.005772 |
| **i-S34 TIN** | 0 | 0 | 0 | 0 | 0 | 0 | 0 | 0 | 0 | 0.010417 | 0 | 0 | 0.003247 |
| **i-S34_S56 TIN** | 0 | 0 | 0 | 0 | 0 | 0 | 0 | 0.005682 | 0.002315 | 0.009549 | 0 | 0 | 0.003517 |
| **i-S34_S56_SGC TIN** | 0 | 0 | 0 | 0 | 0 | 0 | 0 | 0.004941 | 0.016908 | 0.010417 | 0 | 0 | 0.005929 |
| **i-S34_SGC TIN** | 0 | 0 | 0 | 0 | 0 | 0 | 0 | 0 | 0.01634 | 0.006127 | 0 | 0 | 0.010695 |
| **i-S34_SGC_SAC TIN** | 0 | 0 | 0 | 0 | 0 | 0 | 0 | 0 | 0.055556 | 0.010417 | 0 | 0 | 0.016234 |
| **i-S56 TIN** | 0 | 0 | 0 | 0 | 0 | 0 | 0 | 0 | 0 | 0.01 | 0 | 0 | 0.004156 |
| **i-S56_SGC TIN** | 0 | 0 | 0 | 0 | 0 | 0 | 0 | 0.00947 | 0.017361 | 0.00651 | 0 | 0 | 0.006223 |
| **i-S56_SGC_SAC TIN** | 0 | 0 | 0 | 0 | 0 | 0 | 0 | 0.030303 | 0.012346 | 0.013889 | 0 | 0 | 0.007215 |
| **i-S56_SAC TIN** | 0 | 0 | 0 | 0 | 0 | 0 | 0 | 0.030303 | 0 | 0.013889 | 0 | 0 | 0.017316 |
| **i-SGC TIN** | 0 | 0 | 0 | 0 | 0 | 0 | 0 | 0 | 0.007937 | 0.008929 | 0 | 0 | 0.010204 |
| **i-SGC_SAC TIN** | 0 | 0 | 0 | 0 | 0 | 0 | 0 | 0.090909 | 0 | 0.013889 | 0 | 0 | 0 |
| **i-ns TIN** | 0 | 0 | 0 | 0 | 0 | 0 | 0 | 0.007136 | 0.010239 | 0.007679 | 0 | 0 | 0.006427 |
| **SIN** | 0 | 0 | 0 | 0 | 0 | 0 | 0 | 0 | 0 | 0.020833 | 0 | 0 | 0 |
| **TPN-E** | 0 | 0 | 0 | 0 | 0 | 0 | 0 | 0 | 0 | 0 | 0 | 0 | 0 |
| **TPN-O** | 0 | 0 | 0 | 0 | 0 | 0 | 0 | 0 | 0 | 0 | 0 | 0 | 0 |

**Table S2 (continued) (Part 2 of 4; from e-S12_S56_SGC TIN to e-ns TIN).**

| | e-S12_S56_SGC TIN | e-S12_SGC TIN | e-S34 TIN | e-S34_S56 TIN | e-S34_S56_SGC TIN | e-S34_SGC TIN | e-S34_SGC_SAC TIN | e-S56 TIN | e-S56_SGC TIN | e-S56_SGC_SAC TIN | e-SGC TIN | e-SAC TIN | e-ns TIN |
|---|---|---|---|---|---|---|---|---|---|---|---|---|---|
| **SO RGC** | 0 | 0 | 0 | 0 | 0 | 0 | 0 | 0 | 0 | 0 | 0 | 0 | 0 |
| **S12 RGC** | 0.011952 | 0 | 0 | 0 | 0 | 0.001667 | 0.010714 | 0 | 0 | 0 | 0 | 0 | 0 |
| **S34 RGC** | 0.00051 | 0.005952 | 0 | 0 | 0 | 0.004762 | 0.040816 | 0 | 0.049603 | 0 | 0 | 0 | 0.00156 |
| **S56 RGC** | 0.014881 | 0 | 0 | 0 | 0 | 0.005556 | 0.029762 | 0 | 0.069846 | 0 | 0.005208 | 0 | 0.00817 |
| **SGC RGC** | 0 | 0 | 0 | 0 | 0 | 0 | 0.042857 | 0 | 0.093846 | 0 | 0 | 0 | 0.029412 |
| **SAC RGC** | 0 | 0 | 0 | 0 | 0.020833 | 0.011111 | 0.02381 | 0 | 0.018547 | 0 | 0.006944 | 0 | 0.021786 |
| **e-SO TIN** | 0.035714 | 0 | 0 | 0 | 0 | 0 | 0 | 0 | 0 | 0 | 0 | 0 | 0 |
| **e-SO_S56 TIN** | 0.006494 | 0.060606 | 0 | 0 | 0 | 0 | 0.012987 | 0 | 0 | 0 | 0 | 0 | 0.003565 |
| **e-SO_S56_SGC TIN** | 0.007937 | 0 | 0 | 0 | 0 | 0.007407 | 0.015873 | 0 | 0 | 0 | 0.013889 | 0 | 0.004357 |
| **e-S12 TIN** | 0.002976 | 0 | 0 | 0.005952 | 0 | 0.008333 | 0.011905 | 0 | 0 | 0 | 0 | 0 | 0.001634 |
| **e-S12_S34_S56 TIN** | 0.011905 | 0 | 0 | 0 | 0 | 0.011111 | 0 | 0 | 0 | 0 | 0 | 0 | 0.003268 |
| **e-S12_S34_S56_SGC TIN** | 0 | 0 | 0 | 0 | 0.041667 | 0 | 0.047619 | 0 | 0 | 0 | 0 | 0 | 0.013072 |
| **e-S12_S56 TIN** | 0.016917 | 0.013158 | 0 | 0.003759 | 0.001645 | 0.007018 | 0.015038 | 0 | 0.023543 | 0 | 0.006579 | 0 | 0.008514 |
| **e-S12_S56_SGC TIN** | 0.015306 | 0.011905 | 0 | 0.030612 | 0.008929 | 0.004762 | 0.040816 | 0 | 0.012747 | 0 | 0 | 0 | 0.021008 |
| **e-S12_SGC TIN** | 0.02381 | 0.027778 | 0 | 0 | 0 | 0 | 0 | 0.041667 | 0 | 0 | 0 | 0 | 0 |
| **e-S34 TIN** | 0 | 0.166667 | 0 | 0 | 0 | 0 | 0 | 0 | 0 | 0 | 0 | 0 | 0.019608 |
| **e-S34_S56 TIN** | 0.010204 | 0 | 0 | 0 | 0 | 0 | 0 | 0 | 0 | 0 | 0.017857 | 0 | 0.008403 |

| | e-S12_S56_SGC TIN | e-S12_SGC TIN | e-S34 TIN | e-S34_S56 TIN | e-S34_S56_SGC TIN | e-S34_SGC TIN | e-S34_SGC_SAC TIN | e-S56 TIN | e-S56_SGC TIN | e-S56_SGC_SAC TIN | e-SGC TIN | e-SAC TIN | e-ns TIN |
|---|---|---|---|---|---|---|---|---|---|---|---|---|---|
| **e-S34_S56_SGC TIN** | 0.026786 | 0.020833 | 0 | 0.017857 | 0 | 0.008333 | 0 | 0 | 0.013808 | 0 | 0.015625 | 0 | 0.026961 |
| **e-S34_SGC TIN** | 0.02381 | 0.022222 | 0 | 0.028571 | 0 | 0.008889 | 0.028571 | 0 | 0.015128 | 0 | 0.016667 | 0 | 0.022222 |
| **e-S34_SGC_SAC TIN** | 0.030612 | 0.047619 | 0 | 0 | 0 | 0.009524 | 0.020408 | 0 | 0.015495 | 0 | 0 | 0 | 0.033613 |
| **e-S56 TIN** | 0.017857 | 0 | 0 | 0 | 0 | 0 | 0.107143 | 0 | 0 | 0 | 0.03125 | 0 | 0.004902 |
| **e-S56_SGC TIN** | 0.010989 | 0 | 0 | 0.005495 | 0.004808 | 0.010256 | 0.016484 | 0 | 0.002959 | 0 | 0.019231 | 0 | 0.025083 |
| **e-S56_SGC_SAC TIN** | 0.028571 | 0 | 0 | 0 | 0 | 0 | 0.057143 | 0 | 0 | 0 | 0.025 | 0 | 0.003922 |
| **e-SGC TIN** | 0.017857 | 0 | 0 | 0.089286 | 0 | 0.025 | 0.035714 | 0 | 0 | 0 | 0.03125 | 0 | 0.022059 |
| **e-SAC TIN** | 0 | 0 | 0 | 0 | 0 | 0 | 0 | 0 | 0 | 0 | 0 | 0 | 0 |
| **e-ns TIN** | 0.021008 | 0.009804 | 0 | 0.005602 | 0 | 0.007843 | 0.011204 | 0 | 0.001508 | 0 | 0.002451 | 0 | 0.01115 |
| **i-SO TIN** | 0 | 0 | 0 | 0 | 0 | 0 | 0.014286 | 0 | 0 | 0 | 0.0125 | 0 | 0.001961 |
| **i-SO_S12 TIN** | 0 | 0 | 0 | 0 | 0 | 0 | 0 | 0 | 0 | 0 | 0 | 0 | 0 |
| **i-SO_S12_S56 TIN** | 0.02381 | 0 | 0 | 0 | 0 | 0 | 0 | 0 | 0 | 0 | 0 | 0 | 0 |
| **i-SO_S56 TIN** | 0.011429 | 0.005714 | 0 | 0 | 0 | 0.005333 | 0.028571 | 0 | 0 | 0 | 0.005 | 0 | 0.003922 |
| **i-SO_S56_SGC TIN** | 0.02381 | 0 | 0 | 0.015873 | 0 | 0.014815 | 0 | 0 | 0.012821 | 0 | 0.013889 | 0 | 0.008715 |
| **i-S12 TIN** | 0 | 0.004082 | 0 | 0.004082 | 0 | 0.00381 | 0 | 0 | 0 | 0 | 0 | 0 | 0.001681 |
| **i-S12_S34_S56_SGC TIN** | 0 | 0.071429 | 0 | 0.071429 | 0 | 0 | 0 | 0 | 0 | 0 | 0 | 0 | 0 |
| **i-S12_S34_SGC TIN** | 0 | 0 | 0 | 0.047619 | 0 | 0 | 0.047619 | 0 | 0 | 0 | 0 | 0 | 0 |
| **i-S12_S56 TIN** | 0.003695 | 0.011084 | 0 | 0.002463 | 0 | 0.001149 | 0.022167 | 0 | 0.000995 | 0.001437 | 0.00431 | 0 | 0.002874 |
| **i-S12_S56_SGC TIN** | 0.02521 | 0.016807 | 0 | 0.029412 | 0 | 0.005882 | 0.029412 | 0 | 0 | 0 | 0.011029 | 0 | 0.010957 |

| | e-S12_S56_SGC TIN | e-S12_SGC TIN | e-S34 TIN | e-S34_S56 TIN | e-S34_S56_SGC TIN | e-S34_SGC TIN | e-S34_SGC_SAC TIN | e-S56 TIN | e-S56_SGC TIN | e-S56_SGC_SAC TIN | e-SGC TIN | e-SAC TIN | e-ns TIN |
|---|---|---|---|---|---|---|---|---|---|---|---|---|---|
| **i-S12_SGC TIN** | 0.02381 | 0 | 0 | 0 | 0 | 0 | 0 | 0 | 0 | 0 | 0 | 0 | 0.006536 |
| **i-S34 TIN** | 0 | 0 | 0 | 0 | 0 | 0.016667 | 0.035714 | 0 | 0.009615 | 0 | 0 | 0 | 0 |
| **i-S34_S56 TIN** | 0.004464 | 0.017857 | 0 | 0 | 0 | 0.005556 | 0.029762 | 0 | 0.002404 | 0 | 0 | 0 | 0.006127 |
| **i-S34_S56_SGC TIN** | 0.013199 | 0.009317 | 0 | 0.020186 | 0.004076 | 0.004348 | 0.020186 | 0 | 0.000418 | 0 | 0.016304 | 0 | 0.01428 |
| **i-S34_SGC TIN** | 0.02521 | 0.008403 | 0 | 0.016807 | 0 | 0.005882 | 0.004202 | 0 | 0.002262 | 0 | 0.029412 | 0 | 0.012687 |
| **i-S34_SGC_SAC TIN** | 0.035714 | 0 | 0 | 0 | 0 | 0 | 0.035714 | 0 | 0 | 0 | 0 | 0 | 0.009804 |
| **i-S56 TIN** | 0.014286 | 0.005714 | 0 | 0 | 0 | 0.005333 | 0.005714 | 0 | 0 | 0 | 0 | 0 | 0.009412 |
| **i-S56_SGC TIN** | 0.012649 | 0.008929 | 0 | 0.013393 | 0 | 0.002778 | 0.025298 | 0 | 0.001202 | 0 | 0.014323 | 0 | 0.009804 |
| **i-S56_SGC_SAC TIN** | 0.015873 | 0.02381 | 0 | 0.031746 | 0 | 0 | 0.015873 | 0 | 0 | 0 | 0.006944 | 0 | 0.008715 |
| **i-S56_SAC TIN** | 0.047619 | 0.047619 | 0 | 0 | 0 | 0 | 0 | 0 | 0 | 0 | 0 | 0 | 0.026144 |
| **i-SGC TIN** | 0.02551 | 0.020408 | 0 | 0.030612 | 0 | 0.014286 | 0.020408 | 0 | 0.010989 | 0 | 0.017857 | 0 | 0.016807 |
| **i-SGC_SAC TIN** | 0.02381 | 0.095238 | 0 | 0.047619 | 0 | 0 | 0 | 0 | 0 | 0 | 0 | 0 | 0.026144 |
| **i-ns TIN** | 0.011214 | 0.009264 | 0 | 0.006826 | 0.000427 | 0.002958 | 0.014139 | 0 | 0.001969 | 0 | 0.009812 | 0 | 0.010908 |
| **SIN** | 0 | 0 | 0 | 0 | 0 | 0.004167 | 0.053571 | 0 | 0 | 0 | 0.015625 | 0 | 0 |
| **TPN-E** | 0 | 0 | 0 | 0 | 0 | 0 | 0 | 0 | 0 | 0 | 0 | 0 | 0 |
| **TPN_O** | 0 | 0 | 0 | 0 | 0 | 0 | 0 | 0 | 0 | 0 | 0 | 0 | 0 |

**Table S2 (continued) (Part 3 of 4; from i-SO TIN to i-S34_S56 TIN).**

| | i-SO TIN | i-SO_S12 TIN | i-SO_S12_S56 TIN | i-SO_S56 TIN | i-SO_S56_SGC TIN | i-S12 TIN | i-S12_S34_S56_SGC TIN | i-S12_S34_SGC TIN | i-S12_S56 TIN | i-S12_S56_SGC TIN | i-S12_SGC TIN | i-S34 TIN | i-S34_S56 TIN |
|---|---|---|---|---|---|---|---|---|---|---|---|---|---|
| **SO RGC** | 0.001249 | 0 | 0 | 0 | 0 | 0.014652 | 0 | 0 | 0.01221 | 0.010424 | 0.005698 | 0 | 0 |
| **S12 RGC** | 0 | 0.011458 | 0 | 0 | 0 | 0.020595 | 0 | 0 | 0.029514 | 0.019314 | 0.018981 | 0 | 0 |
| **S34 RGC** | 0 | 0.030357 | 0.002381 | 0 | 0 | 0.014286 | 0 | 0 | 0.021463 | 0 | 0.020635 | 0 | 0 |
| **S56 RGC** | 0 | 0 | 0.027778 | 0 | 0.00463 | 0 | 0 | 0 | 0 | 0 | 0 | 0 | 0.000868 |
| **SGC RGC** | 0.013333 | 0.066667 | 0.022222 | 0 | 0 | 0 | 0 | 0 | 0 | 0 | 0 | 0 | 0 |
| **SAC RGC** | 0.026923 | 0 | 0 | 0 | 0.021368 | 0 | 0 | 0 | 0 | 0.00905 | 0 | 0 | 0 |
| **e-SO TIN** | 0 | 0 | 0 | 0 | 0 | 0 | 0 | 0 | 0 | 0 | 0 | 0 | 0 |
| **e-SO_S56 TIN** | 0.009091 | 0.045455 | 0 | 0.003636 | 0.010101 | 0 | 0 | 0 | 0.010972 | 0.002674 | 0.020202 | 0 | 0 |
| **e-SO_S56_SGC TIN** | 0.022222 | 0 | 0 | 0 | 0 | 0.006349 | 0 | 0 | 0.001916 | 0 | 0.012346 | 0 | 0 |
| **e-S12 TIN** | 0 | 0.020833 | 0.013889 | 0 | 0 | 0.00119 | 0 | 0 | 0.013592 | 0 | 0.032407 | 0 | 0 |
| **e-S12_S34_S56 TIN** | 0 | 0.041667 | 0.055556 | 0 | 0 | 0.009524 | 0 | 0 | 0.001437 | 0.004902 | 0.018519 | 0 | 0 |
| **e-S12_S34_S56_SGC TIN** | 0.025 | 0 | 0 | 0 | 0 | 0 | 0 | 0 | 0.008621 | 0 | 0.027778 | 0 | 0 |
| **e-S12_S56 TIN** | 0 | 0.022727 | 0.038961 | 0.002597 | 0.011544 | 0.00334 | 0 | 0 | 0.00515 | 0.000764 | 0.015873 | 0 | 0.000271 |
| **e-S12_S56_SGC TIN** | 0.014286 | 0 | 0.02381 | 0 | 0 | 0 | 0 | 0 | 0.005542 | 0.002101 | 0.015873 | 0 | 0.002976 |
| **e-S12_SGC TIN** | 0 | 0 | 0 | 0 | 0 | 0 | 0 | 0 | 0.002463 | 0 | 0 | 0 | 0.005952 |
| **e-S34 TIN** | 0 | 0 | 0 | 0 | 0.111111 | 0 | 0 | 0 | 0.008621 | 0 | 0 | 0 | 0 |

| | i-SO TIN | i-SO_S12 TIN | i-SO_S12_S56 TIN | i-SO_S56 TIN | i-SO_S56_SGC TIN | i-S12 TIN | i-S12_S34_S56_SGC TIN | i-S12_S34_SGC TIN | i-S12_S56 TIN | i-S12_S56_SGC TIN | i-S12_SGC TIN | i-S34 TIN | i-S34_S56 TIN |
|---|---|---|---|---|---|---|---|---|---|---|---|---|---|
| **e-S34_S56 TIN** | 0.014286 | 0.035714 | 0 | 0 | 0 | 0.004082 | 0 | 0 | 0.003695 | 0 | 0.015873 | 0 | 0 |
| **e-S34_S56_SGC TIN** | 0.0125 | 0 | 0.083333 | 0 | 0 | 0.007143 | 0 | 0 | 0.005388 | 0 | 0.027778 | 0 | 0 |
| **e-S34_SGC TIN** | 0.006667 | 0.05 | 0.066667 | 0.005333 | 0.007407 | 0.011429 | 0 | 0 | 0.005172 | 0 | 0.02963 | 0 | 0.001389 |
| **e-S34_SGC_SAC TIN** | 0 | 0.035714 | 0.047619 | 0.005714 | 0.015873 | 0.004082 | 0 | 0 | 0.006158 | 0.004202 | 0.015873 | 0 | 0.002976 |
| **e-S56 TIN** | 0.025 | 0 | 0 | 0.01 | 0 | 0.007143 | 0 | 0 | 0.008621 | 0 | 0 | 0 | 0 |
| **e-S56_SGC TIN** | 0.007692 | 0.009615 | 0.012821 | 0 | 0.004274 | 0.008791 | 0 | 0 | 0.003979 | 0.001131 | 0.008547 | 0 | 0.002404 |
| **e-S56_SGC_SAC TIN** | 0.016667 | 0.041667 | 0 | 0 | 0.037037 | 0.009524 | 0 | 0 | 0.005747 | 0.004902 | 0 | 0 | 0.006944 |
| **e-SGC TIN** | 0.0375 | 0.0625 | 0 | 0 | 0.013889 | 0.010714 | 0 | 0 | 0.010776 | 0.007353 | 0 | 0 | 0 |
| **e-SAC TIN** | 0 | 0 | 0 | 0 | 0 | 0 | 0 | 0 | 0 | 0 | 0 | 0 | 0 |
| **e-ns TIN** | 0.003922 | 0.014706 | 0.052288 | 0.000784 | 0.004357 | 0.001681 | 0 | 0 | 0.003043 | 0.000577 | 0.013072 | 0 | 0.001225 |
| **i-SO TIN** | 0 | 0 | 0 | 0.004 | 0.011111 | 0.005714 | 0 | 0 | 0.002586 | 0 | 0.022222 | 0 | 0 |
| **i-SO_S12 TIN** | 0 | 0 | 0 | 0 | 0 | 0.014286 | 0 | 0 | 0.002155 | 0 | 0 | 0 | 0 |
| **i-SO_S12_S56 TIN** | 0 | 0 | 0 | 0 | 0 | 0 | 0 | 0 | 0 | 0 | 0 | 0 | 0 |
| **i-SO_S56 TIN** | 0.004 | 0.04 | 0.04 | 0.006667 | 0.008889 | 0.002286 | 0 | 0 | 0.006207 | 0 | 0.013333 | 0 | 0.001667 |
| **i-SO_S56_SGC TIN** | 0.022222 | 0 | 0.037037 | 0.004444 | 0.013889 | 0 | 0 | 0 | 0.007663 | 0.003268 | 0 | 0 | 0.00463 |
| **i-S12 TIN** | 0.002857 | 0.021429 | 0 | 0.001143 | 0 | 0.001681 | 0 | 0 | 0.004187 | 0 | 0.022222 | 0 | 0 |
| **i-S12_S34_S56_SGC TIN** | 0.05 | 0 | 0 | 0.02 | 0 | 0 | 0 | 0 | 0.012931 | 0 | 0.055556 | 0 | 0 |
| **i-S12_S34_SGC TIN** | 0 | 0 | 0 | 0 | 0 | 0 | 0 | 0 | 0.011494 | 0 | 0 | 0 | 0 |
| **i-S12_S56 TIN** | 0.005172 | 0.030172 | 0.017241 | 0.003448 | 0.008621 | 0.003941 | 0 | 0 | 0.006297 | 0.001014 | 0.016284 | 0 | 0.001078 |

| | i-SO TIN | i-SO_S12 TIN | i-SO_S12_S56 TIN | i-SO_S56 TIN | i-SO_S56_SGC TIN | i-S12 TIN | i-S12_S34_S56_SGC TIN | i-S12_S34_SGC TIN | i-S12_S56 TIN | i-S12_S56_SGC TIN | i-S12_SGC TIN | i-S34 TIN | i-S34_S56 TIN |
|---|---|---|---|---|---|---|---|---|---|---|---|---|---|
| **i-S12_S56_SGC TIN** | 0.023529 | 0.036765 | 0.039216 | 0.005882 | 0.01634 | 0.008403 | 0 | 0 | 0.009888 | 0.001783 | 0.026144 | 0 | 0.003676 |
| **i-S12_SGC TIN** | 0.011111 | 0 | 0.037037 | 0 | 0 | 0 | 0 | 0 | 0.000958 | 0 | 0 | 0 | 0.002315 |
| **i-S34 TIN** | 0 | 0 | 0.083333 | 0 | 0 | 0 | 0 | 0 | 0.002155 | 0 | 0 | 0 | 0 |
| **i-S34_S56 TIN** | 0.008333 | 0.03125 | 0.041667 | 0.000833 | 0.006944 | 0.005357 | 0 | 0 | 0.00431 | 0 | 0.011574 | 0 | 0.000443 |
| **i-S34_S56_SGC TIN** | 0.007609 | 0.05163 | 0.025362 | 0.004783 | 0.006039 | 0.006211 | 0 | 0 | 0.00834 | 0.001918 | 0.024155 | 0 | 0.002264 |
| **i-S34_SGC TIN** | 0.005882 | 0.095588 | 0.039216 | 0.004706 | 0.003268 | 0.010924 | 0 | 0 | 0.008874 | 0.002595 | 0.01634 | 0 | 0.004902 |
| **i-S34_SGC_SAC TIN** | 0.025 | 0 | 0 | 0 | 0 | 0 | 0 | 0 | 0.006466 | 0 | 0 | 0 | 0.005208 |
| **i-S56 TIN** | 0.02 | 0.01 | 0.066667 | 0.0032 | 0.017778 | 0.006857 | 0 | 0 | 0.004828 | 0 | 0.008889 | 0 | 0.000833 |
| **i-S56_SGC TIN** | 0.017708 | 0.041667 | 0.013889 | 0.004167 | 0.005787 | 0.005655 | 0 | 0 | 0.007992 | 0.002757 | 0.015046 | 0 | 0.003472 |
| **i-S56_SGC_SAC TIN** | 0.044444 | 0.013889 | 0.018519 | 0.002222 | 0.012346 | 0.003175 | 0 | 0 | 0.009579 | 0.001634 | 0.024691 | 0 | 0.00463 |
| **i-S56_SAC TIN** | 0 | 0 | 0 | 0 | 0 | 0 | 0 | 0 | 0.011494 | 0 | 0.037037 | 0 | 0 |
| **i-SGC TIN** | 0.035714 | 0 | 0.02381 | 0 | 0.007937 | 0.012245 | 0 | 0 | 0.004926 | 0.004202 | 0.02381 | 0 | 0.001488 |
| **i-SGC_SAC TIN** | 0.033333 | 0 | 0 | 0 | 0.037037 | 0 | 0 | 0 | 0.011494 | 0.009804 | 0.037037 | 0 | 0 |
| **i-ns TIN** | 0.009898 | 0.030717 | 0.029579 | 0.004505 | 0.00986 | 0.003803 | 0 | 0 | 0.005914 | 0.001305 | 0.020099 | 0.001706 | 0.001849 |
| **SIN** | 0 | 0.125 | 0 | 0.005 | 0 | 0.0125 | 0 | 0 | 0.010237 | 0 | 0.048611 | 0 | 0 |
| **TPN-E** | 0 | 0 | 0 | 0 | 0 | 0 | 0 | 0 | 0 | 0 | 0 | 0 | 0 |
| **TPN-O** | 0 | 0 | 0 | 0 | 0 | 0 | 0 | 0 | 0 | 0 | 0 | 0 | 0 |

**Table S2 (continued) (Part 4 of 4; from i-S34_S56_SGC TIN to TPN-O)**

| | i-S34_S56_SGC TIN | i-S34_SGC TIN | i-S34_SGC_SAC TIN | i-S56 TIN | i-S56_SGC TIN | i-S56_SGC_SAC TIN | i-S56_SAC TIN | i-SGC TIN | i-SGC_SAC TIN | i-ns TIN | SIN | TPN-E | TPN-O |
|---|---|---|---|---|---|---|---|---|---|---|---|---|---|
| **SO RGC** | 0 | 0 | 0 | 0 | 0 | 0 | 0 | 0 | 0 | 0 | 0.011218 | 0.008282 | 0.152709 |
| **S12 RGC** | 0.001947 | 0 | 0.033333 | 0 | 0 | 0 | 0 | 0 | 0 | 0 | 0.033958 | 0.039406 | 0.125403 |
| **S34 RGC** | 0.00396 | 0 | 0.048214 | 0.002571 | 0.000595 | 0 | 0 | 0 | 0 | 0 | 0.036786 | 0.104702 | 0.031705 |
| **S56 RGC** | 0.004529 | 0 | 0.052083 | 0.001667 | 0.00217 | 0 | 0 | 0 | 0 | 0.000284 | 0.028646 | 0.141375 | 0.028118 |
| **SGC RGC** | 0.008696 | 0.001961 | 0.066667 | 0.002667 | 0.011806 | 0.007407 | 0 | 0 | 0 | 0.000683 | 0.008333 | 0.120508 | 0.107527 |
| **SAC RGC** | 0.014214 | 0.004525 | 0.038462 | 0 | 0.012821 | 0.07265 | 0.115385 | 0.002747 | 0 | 0.004201 | 0 | 0.10901 | 0.085484 |
| **e-SO TIN** | 0 | 0 | 0 | 0 | 0 | 0 | 0 | 0 | 0 | 0 | 0.03125 | 0.011211 | 0.032258 |
| **e-SO_S56 TIN** | 0.007905 | 0 | 0 | 0 | 0.001894 | 0.010101 | 0.030303 | 0 | 0 | 0.001241 | 0.022727 | 0.095801 | 0.080645 |
| **e-SO_S56_SGC TIN** | 0.006039 | 0 | 0 | 0 | 0.003472 | 0 | 0 | 0 | 0 | 0.001517 | 0.027778 | 0.071251 | 0.096774 |
| **e-S12 TIN** | 0.003623 | 0 | 0.03125 | 0.001667 | 0.003038 | 0.00463 | 0 | 0 | 0 | 0.00128 | 0.028646 | 0.049978 | 0.151237 |
| **e-S12_S34_S56 TIN** | 0.003623 | 0 | 0.041667 | 0.013333 | 0.005208 | 0.009259 | 0 | 0 | 0 | 0.001138 | 0.03125 | 0.078475 | 0.08871 |
| **e-S12_S34_S56_SGC TIN** | 0.013587 | 0 | 0 | 0 | 0.002604 | 0 | 0 | 0 | 0 | 0.00256 | 0.015625 | 0.117713 | 0.112903 |
| **e-S12_S56 TIN** | 0.005082 | 0.000382 | 0.045455 | 0.008312 | 0.003923 | 0.007215 | 0.008658 | 0 | 0 | 0.000975 | 0.025974 | 0.105127 | 0.069652 |
| **e-S12_S56_SGC TIN** | 0.009317 | 0 | 0.035714 | 0.002857 | 0.006696 | 0.019841 | 0.047619 | 0 | 0 | 0.000731 | 0.008929 | 0.139654 | 0.114055 |
| **e-S12_SGC TIN** | 0.001553 | 0 | 0 | 0.011429 | 0.005952 | 0.015873 | 0 | 0 | 0 | 0.000975 | 0 | 0.047406 | 0.043779 |
| **e-S34 TIN** | 0.01087 | 0 | 0.25 | 0.04 | 0 | 0 | 0 | 0 | 0 | 0 | 0 | 0.156951 | 0.129032 |
| **e-S34_S56 TIN** | 0.001553 | 0 | 0.035714 | 0 | 0.004464 | 0.015873 | 0 | 0 | 0 | 0.000975 | 0.026786 | 0.080717 | 0.069124 |

| | i-S34_S56_SGC TIN | i-S34_SGC TIN | i-S34_SGC_SAC TIN | i-S56 TIN | i-S56_SGC TIN | i-S56_SGC_SAC TIN | i-S56_SAC TIN | i-SGC TIN | i-SGC_SAC TIN | i-ns TIN | SIN | TPN-E | TPN-O |
|---|---|---|---|---|---|---|---|---|---|---|---|---|---|
| **e-S34_S56_SGC TIN** | 0.01087 | 0 | 0.0625 | 0.02 | 0.009115 | 0.013889 | 0 | 0 | 0 | 0.00128 | 0.007812 | 0.110426 | 0.114919 |
| **e-S34_SGC TIN** | 0.00942 | 0.001961 | 0.033333 | 0.010667 | 0.006944 | 0.014815 | 0 | 0.004762 | 0 | 0.001593 | 0.025 | 0.109716 | 0.137634 |
| **e-S34_SGC_SAC TIN** | 0.017081 | 0 | 0.142857 | 0.005714 | 0.011905 | 0.015873 | 0.047619 | 0 | 0 | 0.003413 | 0.008929 | 0.142217 | 0.119816 |
| **e-S56 TIN** | 0.005435 | 0 | 0 | 0 | 0.007812 | 0 | 0 | 0 | 0 | 0 | 0.03125 | 0.10426 | 0.112903 |
| **e-S56_SGC TIN** | 0.006271 | 0.002262 | 0.009615 | 0 | 0.007612 | 0.008547 | 0 | 0 | 0 | 0.002363 | 0.019231 | 0.074164 | 0.111663 |
| **e-S56_SGC_SAC TIN** | 0.009058 | 0.004902 | 0.041667 | 0.006667 | 0.003472 | 0.018519 | 0 | 0 | 0 | 0.001706 | 0.010417 | 0.066517 | 0.091398 |
| **e-SGC TIN** | 0.016304 | 0 | 0.0625 | 0 | 0.015625 | 0.055556 | 0.083333 | 0 | 0 | 0.002986 | 0 | 0.209081 | 0.185484 |
| **e-SAC TIN** | 0 | 0 | 0 | 0 | 0 | 0 | 0 | 0 | 0 | 0 | 0 | 0.017937 | 0 |
| **e-ns TIN** | 0.005115 | 0.00173 | 0.053922 | 0.006275 | 0.005719 | 0.013072 | 0.006536 | 0 | 0 | 0.001539 | 0.019608 | 0.071661 | 0.091082 |
| **i-SO TIN** | 0 | 0 | 0.025 | 0.004 | 0.002083 | 0.005556 | 0 | 0 | 0 | 0 | 0.01875 | 0.037265 | 0.102258 |
| **i-SO_S12 TIN** | 0.002717 | 0 | 0 | 0.01 | 0.002604 | 0 | 0 | 0 | 0 | 0.000853 | 0.078125 | 0.030269 | 0.086613 |
| **i-SO_S12_S56 TIN** | 0.003623 | 0 | 0.083333 | 0.013333 | 0.006944 | 0 | 0 | 0 | 0 | 0.001138 | 0 | 0.046338 | 0.048387 |
| **i-SO_S56 TIN** | 0.005652 | 0 | 0.04 | 0.0064 | 0.001667 | 0.006667 | 0 | 0 | 0 | 0.000683 | 0.0325 | 0.065112 | 0.082581 |
| **i-SO_S56_SGC TIN** | 0.008454 | 0 | 0.083333 | 0.013333 | 0.006944 | 0.037037 | 0.074074 | 0 | 0 | 0.001517 | 0.013889 | 0.09417 | 0.111111 |
| **i-S12 TIN** | 0.003727 | 0.00084 | 0.042857 | 0 | 0.001488 | 0.006349 | 0.009524 | 0 | 0 | 0.000585 | 0.017857 | 0.058155 | 0.138157 |
| **i-S12_S34_S56_SGC TIN** | 0.021739 | 0 | 0 | 0 | 0.010417 | 0.055556 | 0 | 0 | 0 | 0.003413 | 0.0625 | 0.143498 | 0.096774 |
| **i-S12_S34_SGC TIN** | 0.01087 | 0 | 0 | 0 | 0.003472 | 0 | 0 | 0 | 0 | 0.002275 | 0.083333 | 0.076233 | 0.107527 |
| **i-S12_S56 TIN** | 0.006184 | 0.001014 | 0.023707 | 0.006897 | 0.00449 | 0.009579 | 0.017241 | 0 | 0 | 0.000647 | 0.034483 | 0.285035 | 0.057725 |
| **i-S12_S56_SGC TIN** | 0.013747 | 0.000865 | 0.058824 | 0.005882 | 0.011029 | 0.029412 | 0.019608 | 0.002101 | 0 | 0.003513 | 0.027574 | 0.15996 | 0.052751 |

| | i-S34_S56_SGC TIN | i-S34_SGC TIN | i-S34_SGC_SAC TIN | i-S56 TIN | i-S56_SGC TIN | i-S56_SGC_SAC TIN | i-S56_SAC TIN | i-SGC TIN | i-SGC_SAC TIN | i-ns TIN | SIN | TPN-E | TPN-O |
|---|---|---|---|---|---|---|---|---|---|---|---|---|---|
| **i-S12_SGC TIN** | 0.002415 | 0 | 0.027778 | 0.008889 | 0.006944 | 0.012346 | 0 | 0 | 0.037037 | 0.000379 | 0.020833 | 0.106697 | 0.040932 |
| **i-S34 TIN** | 0.01087 | 0 | 0.0625 | 0.01 | 0 | 0 | 0 | 0 | 0 | 0.00256 | 0.03125 | 0.07287 | 0.080645 |
| **i-S34_S56 TIN** | 0.005435 | 0.001225 | 0.036458 | 0.0075 | 0.003255 | 0.018519 | 0 | 0 | 0 | 0.000498 | 0.023438 | 0.078288 | 0.086022 |
| **i-S34_S56_SGC TIN** | 0.010989 | 0.001598 | 0.038043 | 0.004783 | 0.009624 | 0.016304 | 0.021739 | 0 | 0 | 0.002597 | 0.023777 | 0.129167 | 0.146914 |
| **i-S34_SGC TIN** | 0.012148 | 0.001783 | 0.088235 | 0.003529 | 0.01348 | 0.021242 | 0 | 0.004202 | 0.009804 | 0.002811 | 0.023897 | 0.114614 | 0.15797 |
| **i-S34_SGC_SAC TIN** | 0.024457 | 0 | 0 | 0 | 0.010417 | 0.055556 | 0 | 0.017857 | 0 | 0.004266 | 0.015625 | 0.149103 | 0.107097 |
| **i-S56 TIN** | 0.005652 | 0.001176 | 0.05 | 0.013333 | 0.004167 | 0.006667 | 0 | 0 | 0 | 0.000546 | 0.0075 | 0.102691 | 0.077419 |
| **i-S56_SGC TIN** | 0.011662 | 0.000919 | 0.052083 | 0.005 | 0.00943 | 0.019097 | 0.017361 | 0.000744 | 0 | 0.001991 | 0.016276 | 0.115471 | 0.12668 |
| **i-S56_SGC_SAC TIN** | 0.013889 | 0.001634 | 0.055556 | 0.004444 | 0.014468 | 0.042484 | 0.037037 | 0 | 0 | 0.003982 | 0.020833 | 0.148979 | 0.144265 |
| **i-S56_SAC TIN** | 0.025362 | 0 | 0.083333 | 0.013333 | 0.006944 | 0.074074 | 0.166667 | 0 | 0 | 0.005688 | 0 | 0.22571 | 0.188172 |
| **i-SGC TIN** | 0.015528 | 0.010504 | 0.053571 | 0.008571 | 0.011161 | 0.031746 | 0.047619 | 0 | 0 | 0.002925 | 0.004464 | 0.132928 | 0.14977 |
| **i-SGC_SAC TIN** | 0.01087 | 0 | 0.083333 | 0.026667 | 0.024306 | 0.055556 | 0.111111 | 0 | 0 | 0.005688 | 0 | 0.210762 | 0.123656 |
| **i-ns TIN** | 0.0092 | 0.000803 | 0.049488 | 0.005324 | 0.007608 | 0.016875 | 0.020478 | 0.000731 | 0 | 0.001823 | 0.021758 | 0.116215 | 0.138408 |
| **SIN** | 0.008152 | 0 | 0.015625 | 0 | 0.004557 | 0 | 0 | 0 | 0 | 0.000853 | 0.0625 | 0.10333 | 0.146169 |
| **TPN-E** | 0 | 0 | 0 | 0 | 0 | 0 | 0 | 0 | 0 | 0 | 0 | 0 | 0 |
| **TPN-O** | 0 | 0 | 0 | 0 | 0 | 0 | 0 | 0 | 0 | 0 | 0 | 0 | 0 |

**Table S3. The connection probability of DL_5-HT and SL_5-HT neurons with TINs and TPNs, related to Figure 4.**

| | **DL_5-HT neurons** | **SL_5-HT neurons** |
|---|---|---|
| **e-ns TIN** | 0.038583 | 0 |
| **e-SO TIN** | 0 | 0 |
| **e-SO_S56 TIN** | 0.029326 | 0 |
| **e-SO_S56_SGC TIN** | 0.060932 | 0 |
| **e-S12 TIN** | 0.013441 | 0 |
| **e-S12_S34_S56 TIN** | 0 | 0 |
| **e-S12_S34_S56_SGC TIN** | 0 | 0 |
| **e-S12_S56 TIN** | 0.014007 | 0.00263 |
| **e-S12_S56_SGC TIN** | 0.039171 | 0.01429 |
| **e-S12_SGC TIN** | 0.048387 | 0 |
| **e-S34 TIN** | 0 | 0 |
| **e-S34_S56 TIN** | 0.004608 | 0 |
| **e-S34_S56_SGC TIN** | 0 | 0 |
| **e-S34_SGC TIN** | 0.004301 | 0 |
| **e-S34_SGC_SAC TIN** | 0.043733 | 0 |
| **e-S56 TIN** | 0 | 0 |
| **e-S56_SGC TIN** | 0 | 0 |
| **e-S56_SGC_SAC TIN** | 0.001241 | 0 |
| **e-SGC TIN** | 0 | 0 |
| **e-SAC TIN** | 0.032258 | 0 |

| | DL_5-HT neurons | SL_5-HT neurons |
|---|---|---|
| **i-ns TIN** | 0.044734 | 0 |
| **i-SO TIN** | 0.025806 | 0 |
| **i-SO_S12 TIN** | 0.056452 | 0 |
| **i-SO_S12_S56 TIN** | 0.064516 | 0.04667 |
| **i-SO_S56 TIN** | 0.005161 | 0 |
| **i-SO_S56_SGC TIN** | 0.010753 | 0 |
| **i-S12 TIN** | 0.008295 | 0.02857 |
| **i-S12_S34_S56_SGC TIN** | 0 | 0 |
| **i-S12_S34_SGC TIN** | 0 | 0 |
| **i-S12_S56 TIN** | 0.014461 | 0.04345 |
| **i-S12_S56_SGC TIN** | 0.000949 | 0 |
| **i-S12_SGC TIN** | 0.0681 | 0 |
| **i-S34 TIN** | 0 | 0 |
| **i-S34_S56 TIN** | 0.004032 | 0 |
| **i-S34_S56_SGC TIN** | 0.011921 | 0.00435 |
| **i-S34_SGC TIN** | 0.001898 | 0 |
| **i-S34_SGC_SAC TIN** | 0.177419 | 0.05 |
| **i-S56 TIN** | 0.024516 | 0 |
| **i-S56_SAC TIN** | 0 | 0 |
| **i-S56_SGC TIN** | 0.026546 | 0.00417 |
| **i-S56_SGC_SAC TIN** | 0.023297 | 0 |

| | DL_5-HT neurons | SL_5-HT neurons |
|---|---|---|
| **i-SGC TIN** | 0.002304 | 0 |
| **i-SGC_SAC TIN** | 0 | 0 |
| **SIN** | 0.038306 | 0.0297 |
| **TPN-E** | 0.315536 | 0.049 |
| **TPN-O** | 0.049168 | 0.20226 |
